# Pacific geoduck (*Panopea generosa*) resilience to natural pH variation

**DOI:** 10.1101/432542

**Authors:** Laura H. Spencer, Micah Horwith, Alexander T. Lowe, Yaamini R. Venkataraman, Emma Timmins-Schiffman, Brook L. Nunn, Steven B. Roberts

**Affiliations:** University of Washington, School of Aquatic and Fishery Sciences, 1122 NE Boat St, Seattle, WA 98105, United States; Washington State Department of Natural Resources, 1111 Washington St SE, MS 47027, Olympia, WA 98504, United States; University of Washington, Biological Sciences, 24 Kincaid Hall, Seattle, WA 98105, United States; University of Washington, Genome Sciences, William H. Foege Hall, 3720 15th Ave NE, Seattle, WA 98195, United States

**Keywords:** Aquaculture, comparative physiology, ocean acidification, Panopea generosa, proteomics

## Abstract

Pacific geoduck aquaculture is a growing industry, however, little is known about how geoduck respond to varying environmental conditions, or how the industry will fare under projected climate conditions. To understand how geoduck production may be impacted by low pH associated with ocean acidification, multi-faceted environmental heterogeneity needs to be included to understand species and community responses. In this study, eelgrass habitats and environmental heterogeneity across four estuarine bays were leveraged to examine low pH effects on geoduck under different natural regimes, using targeted proteomics to assess physiology. Juvenile geoduck were deployed in eelgrass and adjacent unvegetated habitats for 30 days while pH, temperature, dissolved oxygen, and salinity were monitored. Across the four bays, pH was lower in unvegetated habitats compared to eelgrass habitats. However this did not impact geoduck growth, survival, or proteomic abundance patterns in gill tissue. Temperature and dissolved oxygen differences across all locations corresponded to differences in growth and targeted protein abundance patterns. Specifically, three protein abundance levels (trifunctional-enzyme β-subunit, puromycin-sensitive aminopeptidase, and heat shock protein 90-α) and shell growth positively correlated with dissolved oxygen variability and inversely correlated with mean temperature. These results demonstrate that geoduck may be resilient to low pH in a natural setting, but other abiotic factors (i.e. temperature, dissolved oxygen variability) may have a greater influence on geoduck physiology. In addition this study contributes to the understanding of how eelgrass patches influences water chemistry.

## Introduction

The Pacific geoduck, *Panopea generosa*, is native to the North American Pacific Coast and is a burgeoning aquaculture species with strong overseas demand as a luxury commodity (Coan et al. 2000; Shamshak and King 2015; Vadopalas et al. 2010). As the largest burrowing clam in the world, cultured geoduck reach upwards of 180mm and are harvested after growing approximately 6-7 years in sub- or intertidal sediment (Vadopalas et al. 2015; Washington DNR website 2017; Washington Sea Grant 2013). The long grow-out period and high per-animal value highlights the importance of site selection for farmers to maximize investment; however, there remains a paucity of data on the optimal environmental conditions for geoduck aquaculture.

As marine calcifiers, geoduck may be vulnerable to ocean acidification due to their reliance on calcite and aragonite (forms of calcium carbonate) for shell secretion (Orr et al. 2005; Weiss et al. 2002), both of which become less biologically available as seawater pH declines with pCO_2_ enrichment (Feely et al. 2008). While there are no ocean acidification studies on *Panopea* clams to date, a growing body of research on marine calcifiers generally indicates that projected low pH will shift organisms’ physiology to the detriment of species-wide abundances and distributions (Pörtner 2008; Pörtner and Farrell 2008). However, broad generalizations of how ocean acidification affects calcifiers are few due to varying pH sensitivity between taxa (Gazeau et al. 2007; Ries et al. 2009) and life stage (Kurihara 2008; Kroeker et al. 2010). For example, in the deeply studied oyster genus *Crassostrea*, Miller et al. (2009) found that larvae of two species varied in their response to elevated pCO_2_, as calcification rates were significantly reduced in the Eastern oyster (*C. virginica*), but the Suminoe oyster (*C. ariakesnsis*) showed no negative response. Thus, lessons learned from other bivalve species cannot directly be applied to geoduck.

The effect of low pH on cultured geoduck needs to be explored to help the aquaculture industry make informed site selection, selective breeding, and investment decisions. For practical application, geoduck ocean acidification studies should best replicate the natural environment in which they are grown. Ninety percent of global production occurs in the Puget Sound estuary of Washington State, where environmental drivers vary between subbasin, season, and diurnal cycle (Moore et al. 2008; Shamshak and King 2015). This habitat heterogeneity exposes geoduck to a variety of secondary stressors when outplanted. Similarly, there is substantial evidence that low pH is not occurring in isolation, but rather in conjunction with changes in other environmental drivers such as temperature (meta-analyses: Byrne and Przeslawski 2013; Harvey et al. 2013; Kroeker et al. 2013), dissolved oxygen (Gobler et al. 2014), and salinity (Przeslawski et al. 2015). Thus, single-stressor studies are limited in their predictive capacity of response to broad scale environmental change. For example, an additive, negative effect of elevated pCO_2_ and temperature was observed in juvenile giant fluted clam survival (*Tridacna squamosa*) (Watson et al. 2012). Another consideration is the incorporation of naturally-occurring diurnal pH variability into ocean acidification studies, as variable pH can have differing effects on marine calcifiers compared to persistent low pH (Review, Boyd et al. 2016). Porcelain crabs, for example, exposed to diurnally variable pH and temperature conditions demonstrated significantly slower metabolism than when crabs were exposed to less variability, or to temperature or pH variability alone (Paganini et al. 2014).

To best predict the effect of ocean acidification on geoduck aquaculture, this project deployed geoduck in variable environmental conditions and leveraged the natural pH differences between eelgrass and unvegetated habitats in Washington State estuaries. Ocean acidification studies are increasingly exploiting naturally low pH systems to monitor the environmental heterogeneity alongside test organisms, in hydrothermal vents (Tunnicliffe et al. 2009; Kerrison et al. 2011), shallow CO_2_ seeps (Duquette et al. 2017), coastal upwelling regions and eutrophic estuaries (Howarth et al. 2011; Thomsen et al. 2013). Compared to controlled laboratory studies, these deployment studies can uniquely incorporate natural ranges and daily cycles in air exposure, temperature, pH, dissolved oxygen, salinity, and food availability. For instance, Ringwood and Keppler (2002) deployed hard clams (*Mercenaria mercenaria*) in the Charleston Harbor estuary in South Carolina while collecting physical-chemical parameters. They observed that while salinity was the primary determinant of growth, pH was also important particularly when salinity was low, and when pH dropped below 7.5, a nuanced finding that is more likely to be captured in a natural environment.

Estuaries along the United States Pacific Coast are ideal, natural mesocosms for examining the effect of ocean acidification on commercially vital calcifiers, as they contain dense macroalgae beds (Bulthuis 1995), environmental conditions that vary considerably between subbasins (Banas et al. 2004; Moore et al. 2008), and have rich communities of native and cultured shellfish (Dethier et al. 2006; Miller et al. 2009; Washington Sea Grant 2015). Furthermore, coastal estuaries have already shifted towards lower pH and warmer temperature averages, and are projected to continue along this trend (Abatzoglou et al. 2013; Busch et al. 2013; Doney et al. 2007; Feely et al. 2012, 2010, 2008; Mote and Salathé 2010). The buffering capacity of macroalgae (seagrass meadows, kelp forests) allows for block-designed experiments to examine the effect of pH, while controlling for varying background environments and maintaining diurnal fluctuations (Middelboe and Hansen 2007; Palacios and Zimmerman 2007; Wahl et al. 2018).

In order to better inform geoduck aquaculture practices, we set out to examine how low pH and other natural variation in environmental conditions influence geoduck growth and physiology, using native eelgrass (*Zostera marina*) as a primary determinant of water chemistry. Physiology was evaluated with a unique two-phase proteomics approach using Selected Reaction Monitoring, with targets identified using Data Independent Acquisition, and selected based on prior environmental stress response studies.

Ocean acidification contributes to an elevation of reactive oxygen species in marine invertebrates (Tomanek 2015). Reactive oxygen species (ROS), or free-radicals, result in oxidative stress and in addition to low pH, higher ROS levels are associated with other environmental stressors including temperature, oxygen variability, salinity, and heavy metals (Review, Lushchak, 2011). Upregulation of anti-oxidants such as catalase, peroxiredoxins, and superoxide dismutase (among others) have consistently been observed in bivalves under low pH and heat stress (Tomanek et al. 2011; Matozzo et al. 2013; Hu et al. 2015), and under heavy metal exposure (Giarratano et al. 2014). In addition to ROS response, ocean acidification is thought to elicit a broader and generic molecular stress response in marine bivalves. Notably, the inducible heat shock proteins are associated with response to hypercapnia, in addition to acute heat, inflammation, and heavy metals, as they act as chaperones to recognize and bind to unfolded or improperly folded proteins (Bozaykut et al. 2014). Induction of HSP90, for example, has been universally observed thus far in bivalve species (Fabbri et al. 2008). Metabolic function is also altered under low pH, hypoxia, and salinity stress, generally shifting to anaerobic metabolism to minimize the mitochondrial ROS production associated with aerobic metabolism (Tomanek 2014).

In addition to antioxidant, metabolic, and general stress-response proteins, this study targeted proteins involved in mitotic growth, detoxification, acid-base balance, and ion regulation (Table 2), all quantified simultaneously to characterize the physiological response in the Pacific geoduck under variable pH environments. This novel application demonstrates the advances in proteomic research and the potential it has to improve aquaculture production.

## Methods

### Experimental Design

*Panopea generosa* juveniles (14.0 ± 0.85 mm) were used in this experiment. Animals were of the same cohort, hatched from broodstock harvested from Puget Sound in Washington State, and reared in a commercial facility in Dabob Bay, WA in controlled conditions (18°C, salinity of 30ppt and pH 8.2). Geoduck were out-planted in four bays throughout Western Washington State from June 21 to July 21, 2016: Fidalgo Bay (FB), Port Gamble Bay (PG), and Case Inlet (CI) in Puget Sound, and Willapa Bay (WB) located off the southwest Pacific Coast of Washington (Table 1, Figure 1). All locations were selected based on the criteria that both *Z. marina* eelgrass beds (“eelgrass”), and unvegetated sediment (“unvegetated”) habitats were present. Clams were placed in 10 cm diameter polymerizing vinyl chloride (PVC) pipes buried in sediment with 5 cm exposed; this method replicates aquaculture techniques. Five clams were placed in each of the 3 tubes in both the eelgrass and unvegetated habitat, with a total of 30 clams across 6 tubes per bay. Pipes were covered with a protective mesh exclosure to limit predation. The replicate structures surrounded and were equidistant to a suite of water quality sensors capturing pH (Honeywell Durafet II Electrode, in custom-built housing), salinity (via conductivity, Dataflow Systems Ltd. Odyssey Conductivity and Temperature Logger), dissolved oxygen (Precision Measurement Engineering MiniDOT Logger), and temperature (via dissolved oxygen probes). Sensors were modified for submersible, autonomous data collection, and logged at 10-minute intervals for the duration of the 30-day outplant.

**Figure 1:**
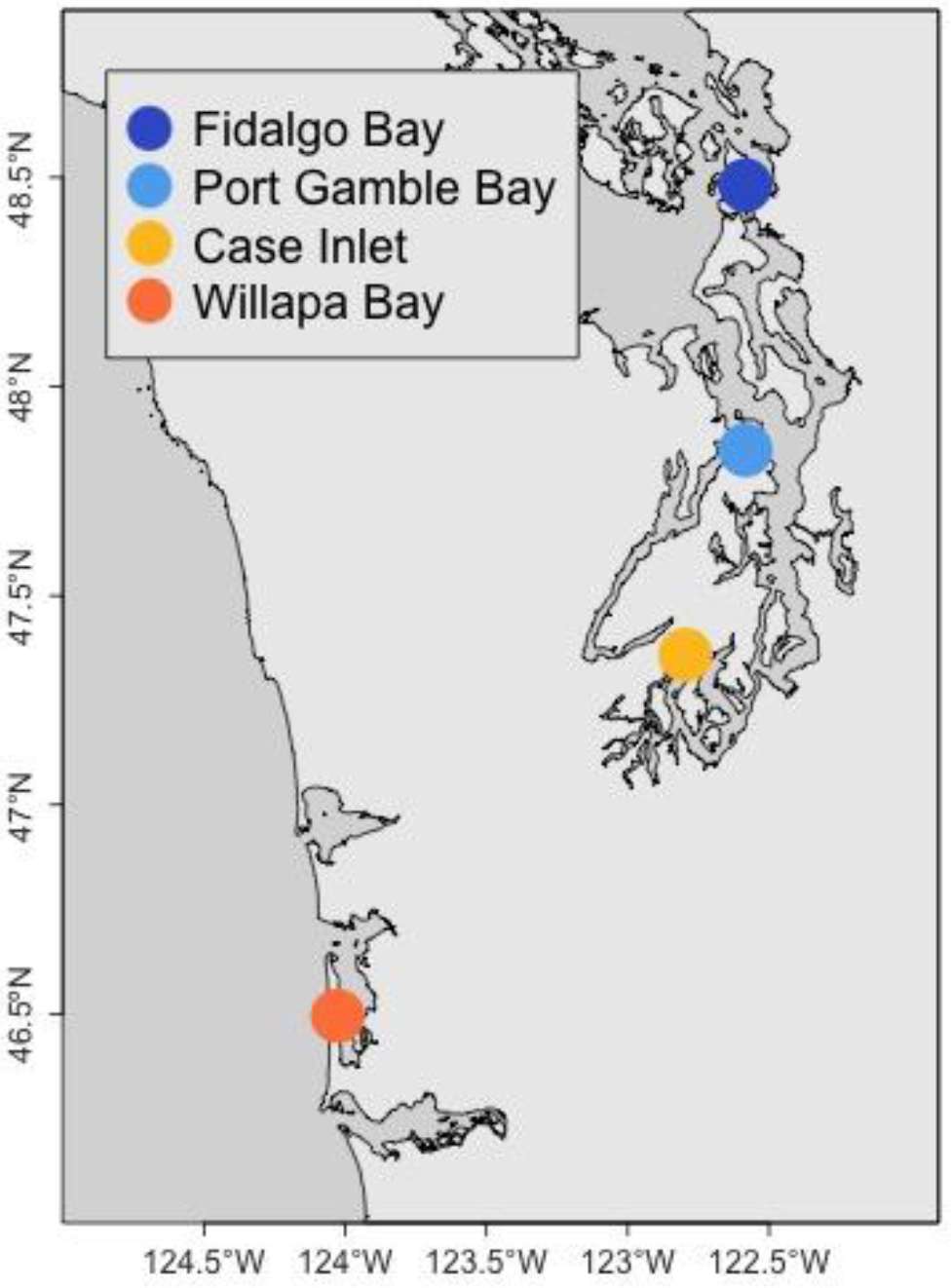
Geoduck juveniles were deployed for 30 days in 2 habitats (eelgrass beds, unvegetated) within 4 bays in Western Washington State

**Table 1:**
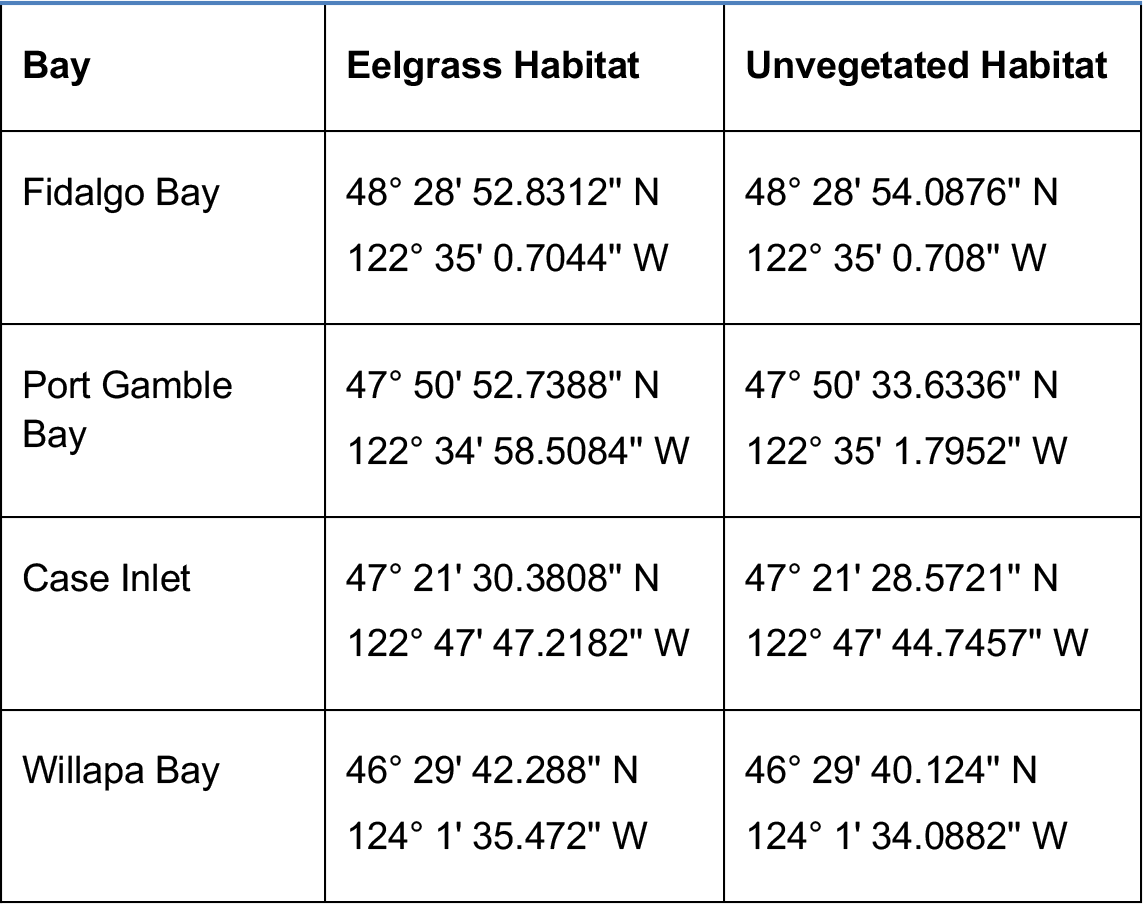
Coordinates for geoduck placement

### Collection and Sampling

Animals were collected during low tide and transferred on wet ice to shore where mortality and size were recorded. Live animals were dissected, and ctenidia tissue was isolated and flash-frozen in an ethanol-dry ice bath. Ctenidia was selected for proteomic analysis due to its direct interaction with the environment, importance in gas and ion regulation, and its implication in environmental stress response (Timmins-Schiffman et al., 2014, Matozzo et al. 2013, Zhang et al. 2015, Thompson et al. 2015). During sampling all instruments were sterilized between samples with bleach then ethanol, and rinsed with nanopure water. Samples were held on dry ice while transported back to the lab and stored at −80ºC.

### Environmental and Growth Data

Temperature, pH, salinity and dissolved oxygen data were compared between outplant locations. Outliers for all environmental parameters were removed, as determined using Tukey Interquartile Range (IQR) method (Tukey 1977), excluding data outside the inner fence (1.5*IQR). Tidal charts from WWW Tide/Current Predictor and salinity data (<20ppt) were also used to remove DO and pH data corresponding to periods of tidal exposure. Four probe failures occurred during deployment (Supplemental Table 1, *failed*) and these data were not included in the analysis (pH at Port Gamble-eelgrass, salinity at Port Gamble-unvegetated & Case Inlet-eelgrass, DO in Fidalgo Bay-eelgrass). Salinity data from two additional locations was not reliably measured (Willapa Bay-unvegetated, Fidalgo Bay-unvegetated), so habitat comparisons were not performed for salinity data. For each parameter at each location, daily mean and daily standard deviation time series were calculated. Relative growth for each animal was determined as (L_f_−L_i_), where L_f_= final geoduck shell length, L_i_ = average initial geoduck shell length within each exclosure (n=5). Differences in growth and environmental parameters between habitat were compared using 2-way analysis of variance (ANOVA) applied to regression models (value ~ habitat*bay). Bays and ad-hoc regions (north vs. south bays) were tested using 1-way ANOVA. Pairwise comparisons were tested with the t-statistic. Significance for all tests was defined as P < 0.05, corrected for multiple comparisons using the Bonferroni correction.

### Protein Analysis

#### Protein Preparation

Relative protein abundance was ultimately assessed in a two-phase proteomics approach using Selected Reaction Monitoring (SRM), with targets identified using Data Independent Acquisition (DIA). Tissues were prepared separately for DIA and SRM, both following the protocol in Timmins-Schiffman et al. (2014) with a few exceptions. For DIA, 8 ctenidia tissue samples were analyzed, one sample from each location and habitat: FB-eelgrass (G048), FB-unvegetated (G058), PGB-eelgrass (G077), PGB-unvegetated (G068), CI-eelgrass (G010), CI-unvegetated (G018), WB-eelgrass (G131), WB-unvegetated (G119). For SRM, new ctenidia samples were examined, 12 individuals per bay (Fidalgo Bay, Port Gamble Bay, Case Inlet, Willapa Bay), with 6 from each habitat (eelgrass, unvegetated) for 48 samples total. Tissue was homogenized with sterile plastic pestle in 100 μl lysis buffer (50 mM NH_4_HCO_3_, 6M urea solution) and sonicated with Sonic Dismembrator (Fisher Scientific, Model 120) at 50% amplitude for ten seconds, three times. Protein concentration was quantified via Pierce™ BCA Protein Assay Kit (ThermoFisher Scientific, Waltham, MA USA).

#### Mini-Trypsin Digestion

Aliquots of protein (30 μg for DIA, 100 μg for SRM) were suspended in Lysis Buffer (50 mM NH_4_HCO_3_ + 6 M urea solution) to a total volume of 100 μl followed by: 1) a 1 hour incubation at 37°C with 200 mM Tris(2-carboxyethyl)phosphine (2.5μl) and 1.5 M Tris at pH 8.8 (6.6 μl); 2) 1 hour at room temperature in dark with 200 mM iodoacetamide (20 μl); 3) 1 hour at room temperature with 200 mM diothiothreitol (20 μl); 4) 1 hour at room temperature with 2 ug/μl Lysyl Endopeptidase (Lys-C, Wako Chemicals) (3.3 μg); 5) overnight at room temperature in 25 mM NH_4_HCO_3_ (800 μl) + high pressure liquid chromatography grade methanol (200 μl) + Pierce Trypsin Protease, MS Grade (1 ug/μl, Thermo Scientific) at 1:30 enzyme:protein ratio (3.3 μg). Samples were evaporated to near dryness at 4°C using a CentriVap Benchtop Vacuum Concentrator.

#### Desalting

Samples were desalted to isolate peptides using MacroSpin Columns (Nest Group, 50-450 μl, Peptide Protein C18). Peptides were reconstituted in 5% acetonitrile + 0.1% trifluoroacetic acid (TFA) (100 μl), then 10% formic acid (70 μl) was added to achieve pH ≤2. Columns were washed with 60% acetonitrile + 0.1% TFA (Solvent A, 200 μl) four times, then equilibrated with 5% acetonitrile + 0.1% TFA (Solvent B, 200 μl) three times. Peptides were bound to the column by running the digest through the column twice, followed by peptide elution with two additions each of Solvent A (100 ul). Columns were spun for 3 minutes at 3000 rpm on VWR Galaxy 16DH digital microcentrifuge at each stage. Samples were evaporated to near dryness at 4°C, then reconstituted in the Final Solvent (3% acetonitrile + 0.1% formic acid) (60 μl for 0.5 μg/μl final concentration of protein, and 50 μl for 2 μg/μl final concentration for DIA & SRM, respectively).

#### Peptide sample preparation and internal standard

Final mixtures for mass spectrometry included 3.13 fmol/μl Peptide Retention Time Calibration mixture (PRTC), 0.33 μg/μl and 0.5 μg/μl peptides for DIA and SRM, respectively, in Final Solvent for 15 μl total volume. To confirm that peptides were quantified correctly in SRM, 10 μg from 5 randomly selected geoduck peptide samples were pooled, and 8 dilutions were prepared by combining with oyster peptides at known percentages of total protein content (10%, 13.3%, 20%, 40%, 60%, 80%, 87.7%, 90%) and analyzed alongside other samples.

### Data Independent Acquisition

#### Data acquisition

Data Independent Acquisition (DIA) was performed to assess global protein abundance patterns and to identify consistently detectable peptides for SRM targets. Eight samples, one per deployment location, were analyzed in technical duplicates via liquid chromatography tandem mass spectrometry (LC-MS/MS) with the Thermo Scientific™ Orbitrap Fusion Lumos™ Tribrid™. Prior to sample analysis, the 30 cm analytical column and 3 cm trap were packed in-house with with C18 beads (Dr. Maisch HPLC, Germany, 0.3 μm). For each sample, 3 μl of geoduck peptides (1.0 μg) + PRTC peptides was injected and analyzed in MS1 over 400-900 *m/z* range, in 5 *m/z* overlapping isolation windows from 450-850 *m/z* with 15K resolution in MS2. Final Solvent blanks were run between each geoduck peptide injection to ensure against peptide carry-over. Lumos MS/MS method and sequence files are available in the project repository (Spencer et al. 2019), and data are available via ProteomeXchange with identifier PXD012266.

#### Protein identification and analysis

Proteins were inferred using an assembled, translated, and annotated *P. generosa* gonad transcriptome (combined male and female) (Timmins-Schiffman et al. 2017; DOI: 10.17605/OSF.IO/3XF6M). Transcriptome peptides were queried against those detected by the Lumos MS/MS using PEptide-Centric Analysis (PECAN) (Ting 2017) to create a peptide spectral library (.blib type file). DIA raw files were first demultiplexed using MSConvert (ProteoWizard version 3.0.10738, 2017-04-12) (Chambers et al. 2012) with filters set to vendor centroiding for msLevels [2,3] (--”peakPicking true 1-2”), and optimizing overlapping spectra (“demultiplex optimization-overlap only”). The transcriptome fasta file was tryptic digested *in silico* in Protein Digestion Simulator (version 2.2.6471.25262), set to Fully Tryptic from 400-6000 fragment mass range, 5 minimum residues allowed, 3 maximum missed cleavages and peak matching thresholds set to 5 ppm mass tolerance, and 0.05 ppm NET tolerance. Skyline version 3.7.0.11317 (MacLean et al. 2010) automatically selected transition peaks and quantified peptide abundances using peak area integration. All PRTC peptide peak selections were manually verified and corrected. Skyline peak selection error rate was calculated by manually checking chromatograms from 100 proteins across all DIA samples. Auto-selected peaks were assigned correct or incorrect selection based on transition retention time alignment across replicates, using PRTC peptides as a reference. Transition peak area, which is assumed to correlate to peptide transition abundance, was exported from Skyline for analysis in R version 2.4-3 (R Core Team 2016). Abundance was normalized by the total ion current (TIC) for each injection. Technical replicate, bay and habitat differences were assessed to inform SRM analysis via non-metric multidimensional scaling (NMDS) analysis using `metaNMDS` in the vegan package (Oksanen et al. 2016) on log(x+1) transformed abundances using a Bray-Curtis dissimilarity matrix. Technical replicate spectral abundances clustered together on NMDS plots, thus were averaged across each sample. Bay and habitat differences in global abundance were visually but not quantitatively analyzed (Supplemental Figure 4).

### Selected Reaction Monitoring

#### Target selection

Thirteen proteins were selected for SRM targets (Table 2). First, candidate proteins (~200) were selected based on biological function listed in the Universal Protein Knowledgebase (Apweiler et al. 2004) and evidence of stress response in bivalves from the scientific literature. Candidate proteins were screened for detectability using DIA results. Selected proteins were required to have ≥3 high quality peptides, each with ≥3 transitions, present in all DIA biological and technical replicate data. Quality peptides had uniform peak morphology and retention time in Skyline across replicates. A total of 49 peptides were selected for SRM: 39 to quantify 13 target proteins (116 transitions), and 10 for internal standard (30 transitions). A full list of transition targets are published on https://panoramaweb.org/e0TsuK.url and available in the project repository (Spencer et al. 2019).

**Table 2:**
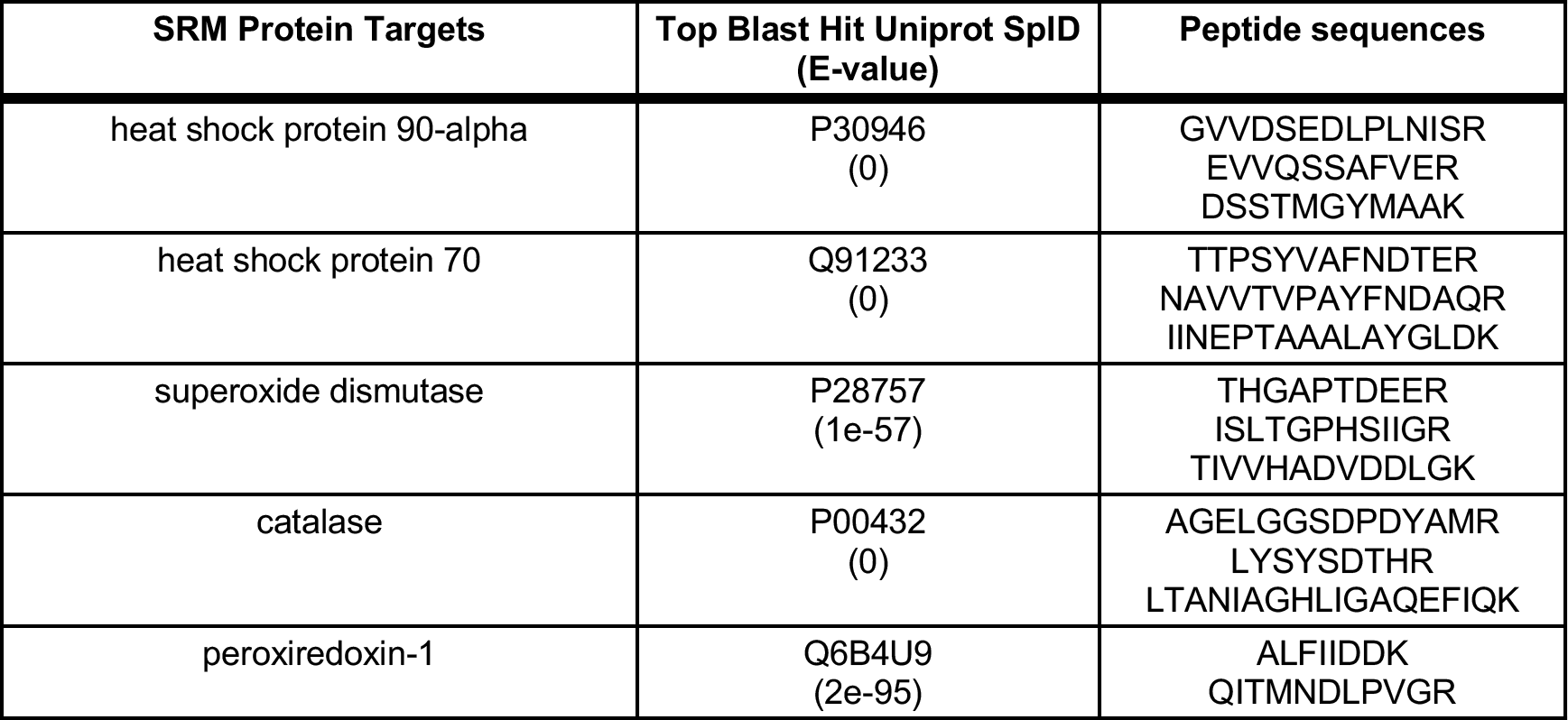
SRM proteins targets selected based on biological function and detectability across DIA samples

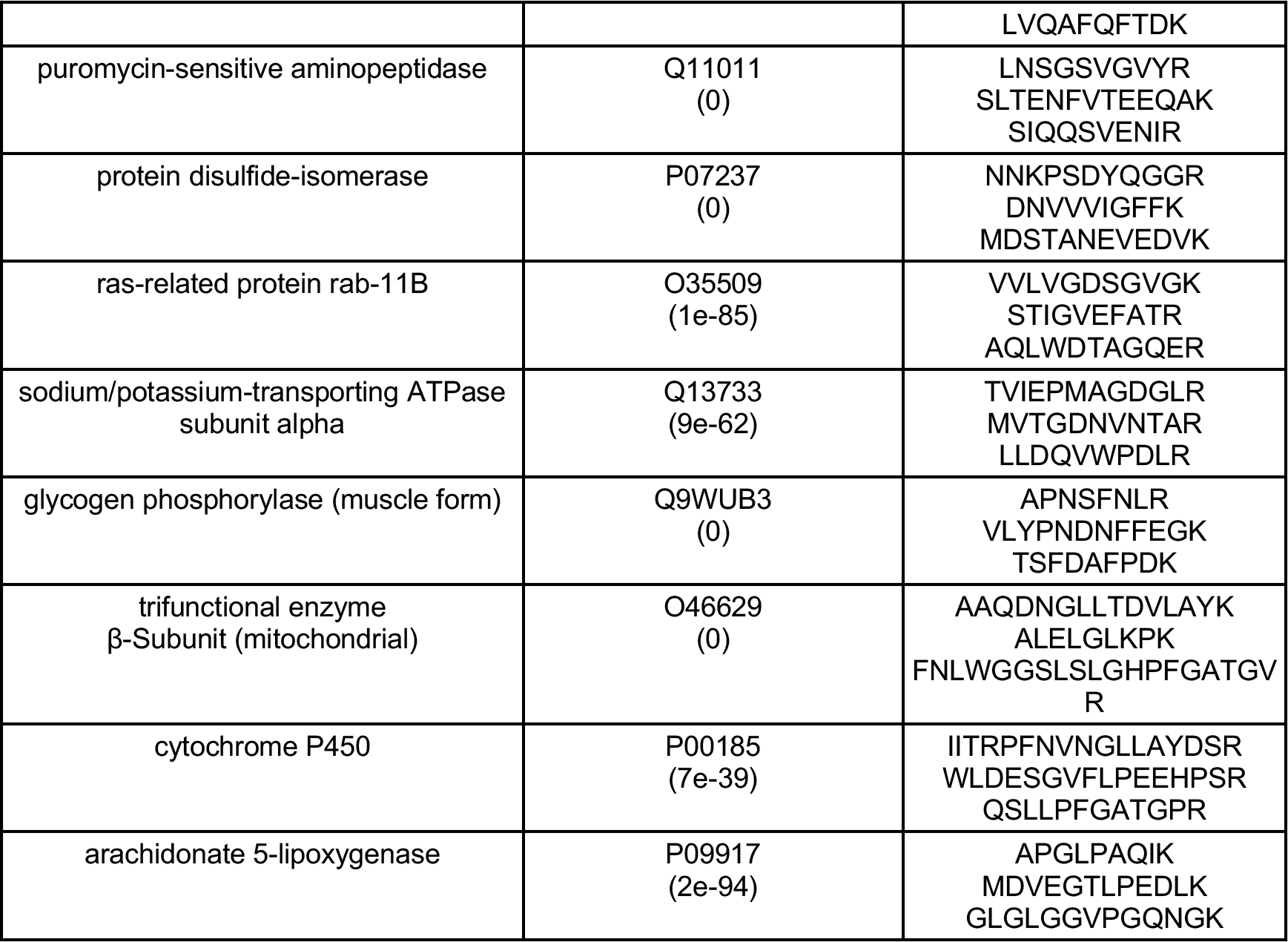

#### Data acquisition

SRM samples were analyzed on a Vantage Triple-Stage Quadrupole Mass Spectrometer (Thermo Scientific, San Jose, CA, USA), and injected by a nanoACQUITY UPLC^®^ system (Waters, Milford, MA, USA) at random in two technical replicates. For each sample, 2 μl of peptides + PRTC solution containing 1.0 μg of geoduck peptides was injected, trapped on a 3 cm pre-column and separated on a 30 cm analytical column using a chromatography gradient of 2-60% acetonitrile over 60 minutes. Columns were prepared as in DIA (above). Samples were injected in randomized groups of 5, followed by a Peptide Retention Time Calibration (PRTC) plus bovine serum albumin peptides (BSA) standard, then Final Solvent blank. Vantage MS sequence and method files are available in the project repository (Spencer et al. 2019).

#### Protein identification and quality assurance

Peptides were identified and quantified via Skyline-daily version 3.7.1.11357 (MacLean et al. 2010). Raw SRM files were imported into a Skyline-daily project along with the target protein transitions, and the spectral library (.blib file) created previously in the DIA Protein Identification step. SRM peptides were verified by regressing PRTC peptide retention time (RT) in SRM against retention time in DIA. A fitted model from PRTC peptides predicted RT of protein target peptides. Where necessary, peak selection and boundaries were manually adjusted for SRM peptide chromatograms, and actual RT were regressed against predict RT to confirm correct selection (F(1,38): 5768, p-value: < 2.2e-16, Adjusted R-squared: 0.9933) (Supplementary Figure 5). Transition peak area, defined henceforth as abundance, was exported from Skyline for further analysis in R (R Core Team 2016). Abundance results from the separate serial dilution samples were used to remove peptides that did not adhere to the dilution curve. Briefly, dilution abundances (exported from Skyline) for each transition were normalized by the most dilute sample abundance, then regressed against predicted ratios. Peptides with slope coefficient 0.2<x<1.5 and adjusted R^2^ >0.7 were included in analysis. Ten of the 39 peptides were discarded from the dataset based on dilution standards results (Supplemental Figure 6). To determine and remove disparate technical replicates, NMDS analysis was performed as described above. Technical replicates with ordination distance >0.2 were removed, and only samples with two technical replicates were preserved for analysis (Supplemental Figure 7). Thirteen technical replicates from different samples and all replicates from three sample were discarded, for 84% technical replicate and 94% biological replicate retention. Within samples, transitions with coefficients of variation (CV) > 40% between technical replicates were also discarded (2% of all transitions across 21 samples). In final dataset for differential analysis, 10 proteins, 26 peptides, and 77 transitions were retained. Mean transition abundance was calculated for replicates, with zero in the place of n/a values, which Skyline generates for replicates without peaks. Transition abundances within each peptide were summed for a total peptide abundance before analyzing for differential abundance.

### Differential protein analysis

After data quality screening, peptide abundance was analyzed for differences between locations and habitats. NMDS plots visualized patterns in peptide abundances by bay and habitat as described above. Global peptide abundance was compared between bay and habitats using two-way ANOVA on log-transformed abundances. For protein-specific comparisons, peptide abundances were grouped by protein, box-cox transformed (Box and Cox 1964) and normality confirmed via qqplot (Wickham 2017). Two-way ANOVA tested abundances for each protein between eelgrass and unvegetated habitats within and between bays. Pairwise comparisons for differentially abundant proteins were tested with the t-statistic. Peptides within proteins were regressed against each other to confirm stable abundance patterns. For all statistical analyses, significance was defined as alpha ≤ 0.05, corrected for multiple comparisons using the Bonferroni correction.

### Correlative analysis

To understand how environmental and biometric parameters covaried, Pearson's product-moment correlation and scatter plots were assessed between protein abundances, growth, and environmental summary statistics (mean and variance). Each protein was assessed independently. Due to salinity probe malfunction, salinity data were not included in correlation tests.

All analyses were performed in RStudio version 1.1.383 (R Core Team 2016). R scripts and notebooks (Spencer et al. 2019), raw data (ProteomeXchange PXD012266), and Skyline project files (https://panoramaweb.org/e0TsuK.url) are publicly available.

## Results

### Environmental & Growth Data

Mean pH differed significantly between habitats across all bays (F(1,206)=180.0, p=1.1e-28) (Figure 2). During the deployment, pH was recorded from 6.71 to 8.34, with mean pH 7.86±0.15 in eelgrass, and 7.51±0.25 in unvegetated habitats (means are for all locations). Variability in pH was significantly different among bays (F(3,206=43.8, p=1.0e-20). Variability did not differ between habitats across all bays, but differences were detected between habitats within Case Inlet and within Willapa Bay (less variable in eelgrass, Supplementary Table 1). The locations with the highest and lowest daily mean pH were Fidalgo Bay-eelgrass (7.90±0.19) and Port Gamble Bay-unvegetated, respectively (7.32±0.25). On average across all locations, pH fluctuated daily by 0.46±0.23 pH units. Considerable heterogeneity among bays was observed in the other environmental parameters. Mean temperature was significantly different among all bays (F(3,236) =129.4, p=2.2e-48), and temperature decreased with latitude (coldest in northernmost Fidalgo Bay, warmest in southernmost Willapa bay). Temperature did not differ between habitats within bays (Supplemental Figure 1). Dissolved oxygen (DO) varied among bays in both daily standard deviation (F(3,210)=132.8, p=4.6e-47) and mean (F(3,210)=56.7, p=1.1e-25). DO variability was substantially higher in the two northern bays (SD was 5.6 and 3.9 mg/L in FB, PGB), as compared to the southern bays (2.5 and 1.4mg/L in CI, WB). Across all bays, DO variability did not differ between habitats, but did differ within Case Inlet and Fidalgo Bay (Supplementary Table 1 & Figure 2). Mean salinity differed by bay (F(3,136)=254.3, p=2.3e-54), with the largest differences between Fidalgo Bay (mean 29.9 ppt) and the other three bays (mean 23.4-27.0 ppt) (Supplemental Figure 3). Growth significantly differed between northern and southern bays (F(1,97)=54.8, P=4.9-11), but not between habitats either within or across all bays. Geoduck in Fidalgo Bay and Port Gamble Bay grew larger compared to Willapa Bay, and Case Inlet (Figure 3). Survival did not differ among locations (Supplementary Table 1).

**Figure 2:**
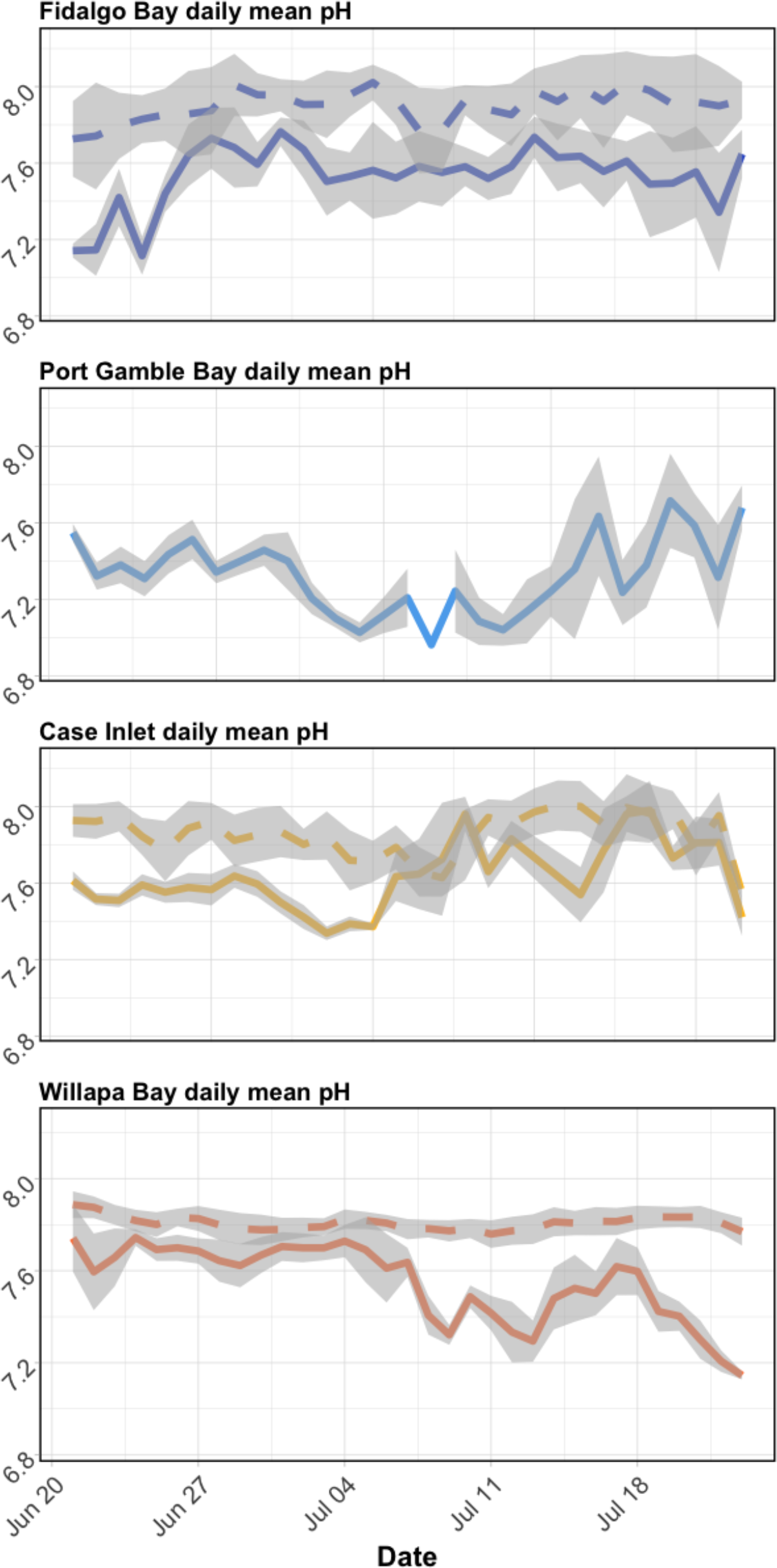
Daily mean pH in eelgrass (dashed lines) and unvegetated (solid lines) across bays during geoduck deployment. Gray ribbons denote daily standard deviation. Data from Port Gamble bay eelgrass are not included due to probe failure.

**Figure 3:**
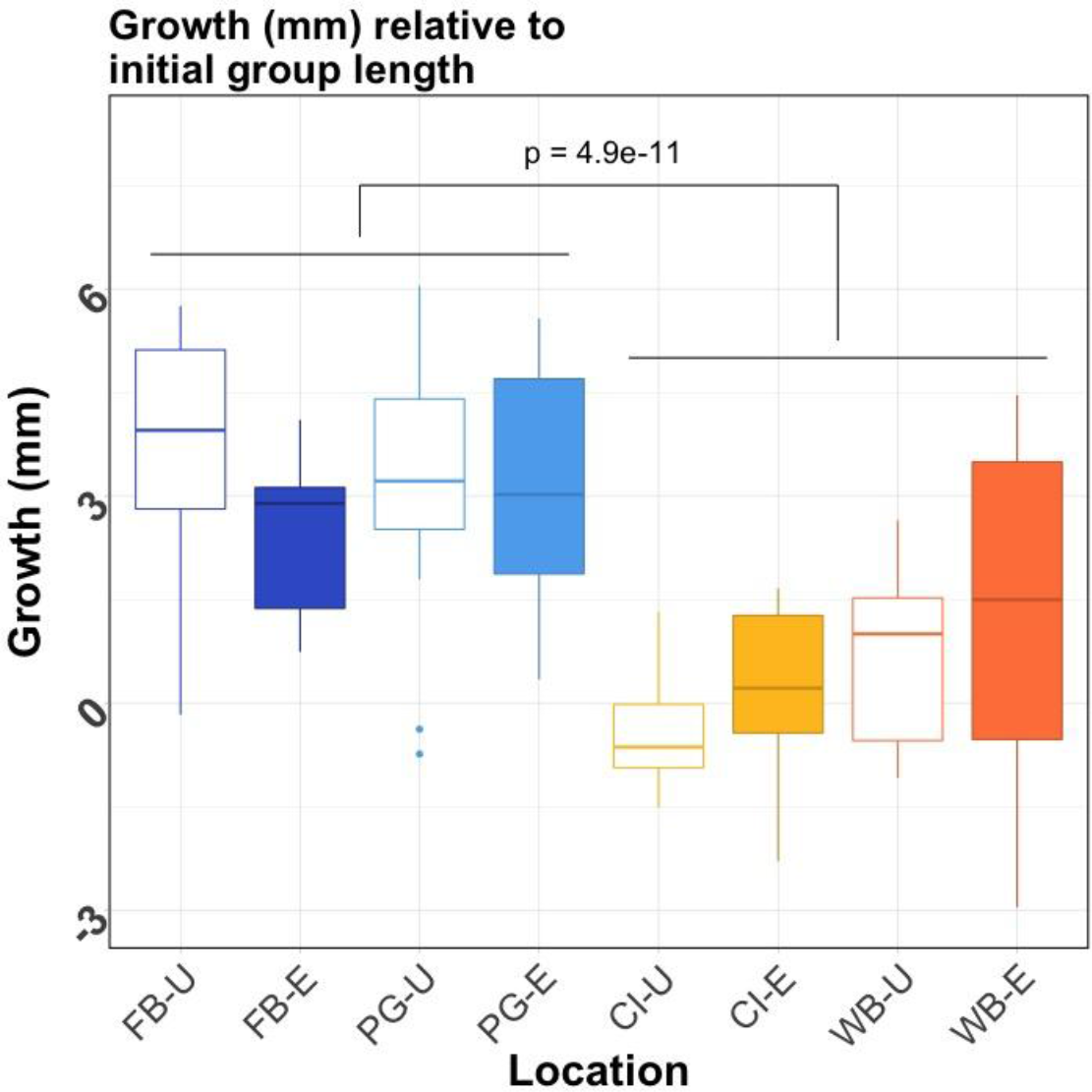
Geoduck shell growth after 30 days across Willapa Bay (WB), Case Inlet (CI), Port Gamble Bay (PG), and Fidalgo Bay (FB), where -U and -E represent unvegetated (empty boxes) and eelgrass habitats (filled boxes), respectively. Growth is relative to the mean initial shell length within deployment groups (n=5 per group, 3 groups per location). Boxes contain all biological replicates lying within the interquartile range (IQR), with median growth indicated by line in middle of boxes. Whiskers extend to the largest value no greater than 1.5*IQR, and dots indicate outliers beyond 1.5*IQR. Geoduck that did not survive deployment are not included. Growth differed significantly between southern bays (WB, CI) and northern bays (PG, FB) (p=4.9e-11) but not between habitats within bays.

### Protein Detection and Variance

In DIA, a total of 298,345 peptide transitions were detected from 30,659 distinct peptides across the 8 samples (one sample per habitat from each bay). These peptides were associated with 8,077 proteins, and more than half of the proteins (4,252) were annotated using Universal Protein Resource database (UniProt). Automated peak selection (Skyline) success rate was 71%.

In SRM, the final dataset after screening included 10 proteins, 26 peptides, and 77 transitions. The 3 proteins fully removed from the dataset were heat shock protein 70, peroxiredoxin-1, and ras-related rab. The SRM mean coefficients of variation (CV) of technical replicate abundances across all transitions decreased from 18.2% to 9.6% after screening. Transition abundance CV within bays ranged from 24.9% to 83.2% with mean 50.1%, and within deployment locations CV ranged from 11.6% to 93.0% with mean 48.9% (Supplemental Table 3). Within proteins, regression analysis indicated that peptide abundances from the same protein differed slightly, however the relative abundances across samples was consistent. This indicates a small degree of background variability in peptide detection, digestion, or stability within proteins that applied to all samples (Pep1xPep2: R^2^_A_=0.985, coefficient=0.682; Pep1xPep3: R^2^_A_=0.990, coefficient=0.954; Pep2xPep3: R^2^_A_=0.990, coefficient=0.954).

### Protein Abundance Differences

None of the 10 targeted proteins were differentially abundant between habitats within or across bays (Figure 4, Supplemental Table 2). NMDS plots of all transitions in DIA and those targeted in SRM revealed clustering of overall proteomic response by bay (Supplemental Figures 4 & 8). In SRM, Fidalgo Bay and Port Gamble Bay samples clustered together (henceforth “northern bays”), and some overlap between Case Inlet and Willapa Bay (“southern bays”) indicated similar protein abundances within these ad-hoc regions (Supplemental Figure 8). This was verified from the ANOVA results, which detected significant abundance differences between northern and southern bays for three proteins: HSP90-α (HSP90) (F(1,133)=20.5, p-adj=1.8e-4), trifunctional-enzyme subunit β-subunit (TEβ) (F(1,88)=11.1, p-adj=0.018), and puromycin-sensitive aminopeptidase (PSA) (F(1,130)=9.11, p-adj=0.043). HSP90 and TEβ abundances were also significantly different between bays (respectively: F(3,131)= 7.80, p-adj=0.0011; F(3,345)= 5.19, p-adj=0.034), but these differences were driven by regional differences, as post-hoc tests detected no differences between Case Inlet and Willapa Bay (southern), or beteen Fidalgo Bay and Port Gamble Bay (northern). For the three differentially abundant proteins, abundances were lowest in Case Inlet (southernmost in Puget Sound) followed by WIllapa Bay (southernmost overall), then Port Gamble Bay, and highest in Fidalgo Bay (northernmost).

**Figure 4:**
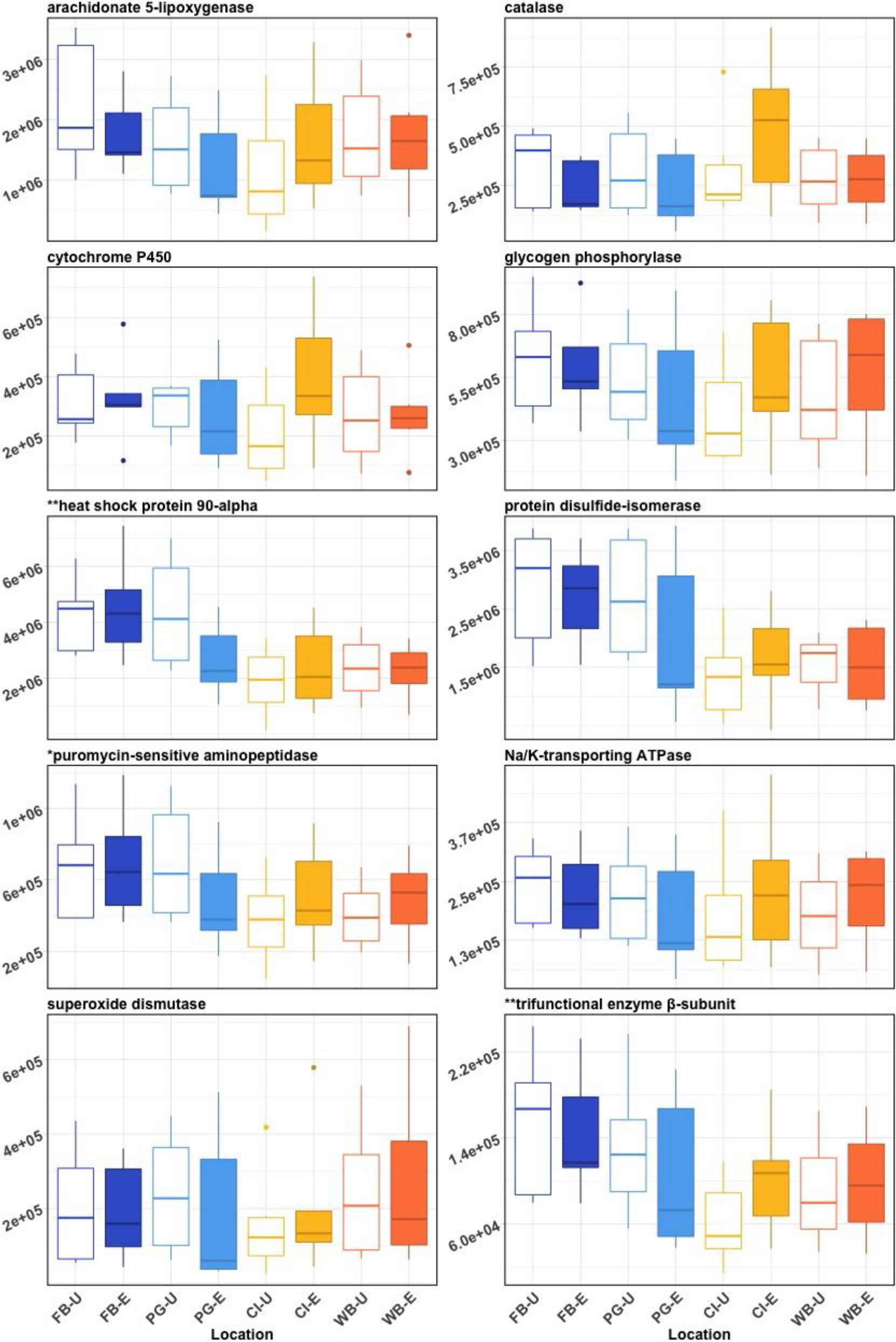
Boxplots of protein mean spectral abundances (mean of 2 or 3 peptides targeted for each protein) for Fidalgo Bay (FB), Port Gamble Bay (PG), Case Inlet (CI), and Willapa Bay (WB), where -U and -E represent unvegetated (white boxes) and eelgrass (filled boxes) habitats, respectively. For each location, n=5 or 6 geoduck. Boxes contain all biological replicates lying within the interquartile range (IQR), with median abundances indicated by line in middle of boxes. Whiskers extend to the largest value no greater than 1.5*IQR, and dots indicate outliers beyond 1.5*IQR. Protein abundance ranges (y-axes) vary between proteins. Differentially abundant proteins between region and bay are indicated by (**), and region only by (*). No protein abundances were significantly different between habitats.

### Correlation between Environment, Abundance, and Growth

Growth positively correlated with peptide abundance in all but 2 of the 10 targeted proteins (no correlation with catalase and superoxide dismutase), including the three proteins that were differentially abundant between bays (Table 3). Growth also correlated with most environmental parameters (excluding salinity SD). Heat Shock Protein 90 correlated positively with dissolved oxygen SD. Mean and SD pH did not correlate with any peptide abundance patterns or growth. None of the environmental parameters, nor growth, correlated significantly with peptide abundances pooled across all proteins.

**Table 3:**
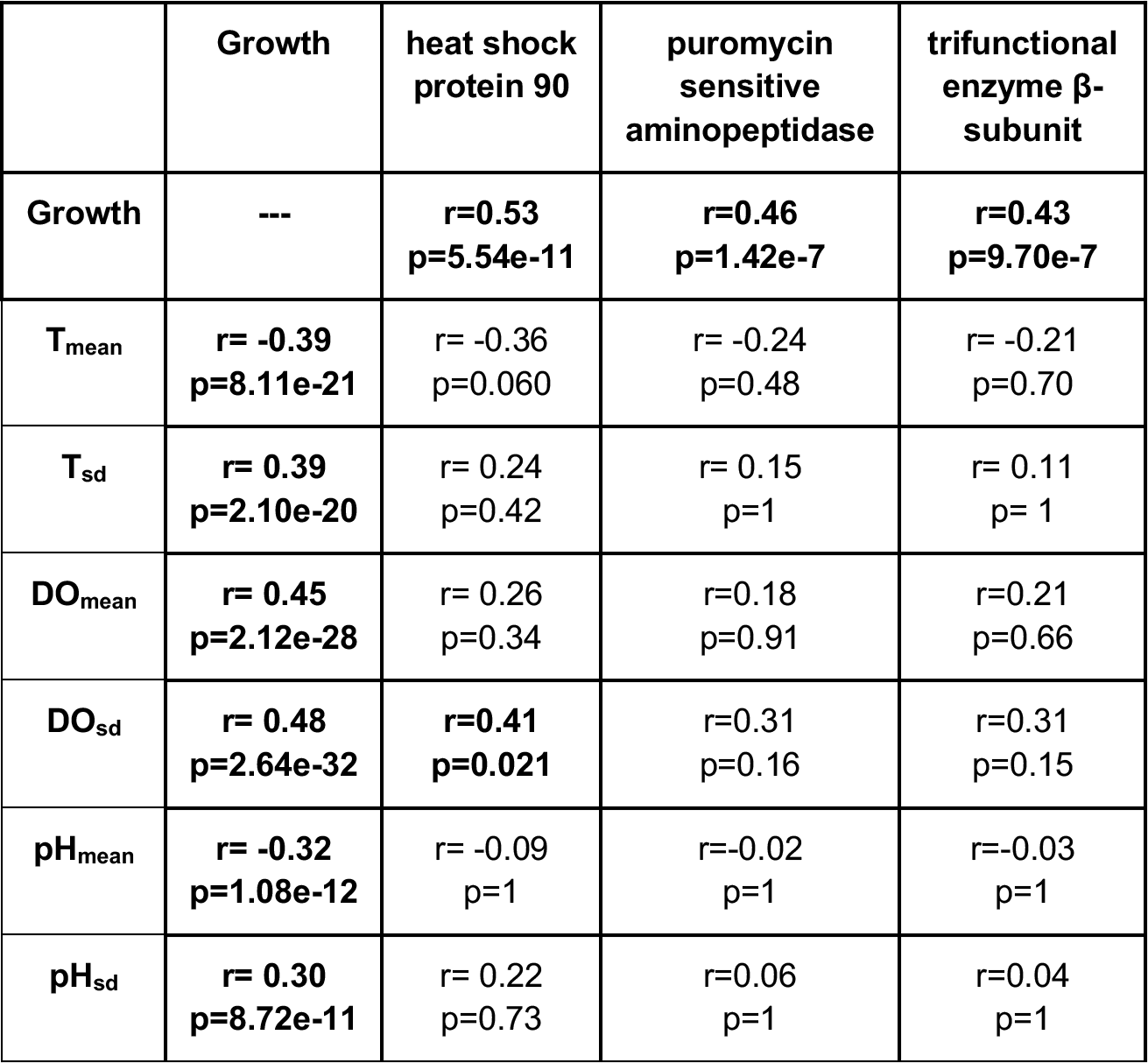
Correlation analysis results between growth, environmental parameters, and peptide abundance (z-transformed). Correlation coefficient r shown with p-values adjusted via Bonferroni correction (number of comparisons). Correlations deemed significant are in bold.

| | Growth | heat shock protein 90 | puromycin sensitive aminopeptidase | trifunctional enzyme β-subunit |
| --- | --- | --- | --- | --- |
| Growth | --- | **r=0.53**<br>**p=5.54e-11** | **r=0.46**<br>**p=1.42e-7** | **r=0.43**<br>**p=9.70e-7** |
| T<sub>mean</sub> | **r= -0.39**<br>**p=8.11e-21** | r= -0.36<br>p=0.060 | r= -0.24<br>p=0.48 | r= -0.21<br>p=0.70 |
| T<sub>sd</sub> | **r= 0.39**<br>**p=2.10e-20** | r= 0.24<br>p=0.42 | r= 0.15<br>p=1 | r= 0.11<br>p=1 |
| DO<sub>mean</sub> | **r= 0.45**<br>**p=2.12e-28** | r= 0.26<br>p=0.34 | r=0.18<br>p=0.91 | r=0.21<br>p=0.66 |
| DO<sub>sd</sub> | **r= 0.48**<br>**p=2.64e-32** | **r=0.41**<br>**p=0.021** | r=0.31<br>p=0.16 | r=0.31<br>p=0.15 |
| pH<sub>mean</sub> | **r= -0.32**<br>**p=1.08e-12** | r= -0.09<br>p=1 | r=-0.02<br>p=1 | r=-0.03<br>p=1 |
| pH<sub>sd</sub> | **r= 0.30**<br>**p=8.72e-11** | r= 0.22<br>p=0.73 | r=0.06<br>p=1 | r=0.04<br>p=1 |

## Discussion

This study tested the effects of varying pH on geoduck, a valuable aquaculture species in a natural setting, and confirmed that *Zostera marina* eelgrass can effectively alter local pH during warm summer months (June and July). We have shown that ocean acidification research on cultured shellfish can augment findings from controlled laboratory studies with field deployments to incorporate natural variability and relevant environmental drivers associated with an organism’s habitat. Targeted proteomics was assessed alongside growth and environmental data for an integrated view of how geoduck respond to varying environmental conditions. Proteomics is a powerful approach suitable for comparative physiological studies of non-model, marine organisms (Tomanek, 2014). Using a two step method, this study detected substantially more proteins (8,077) compared to the previous geoduck study using Data Dependent Acquisition (3,651) (Timmins-Schiffman et al. 2017). This produced a valuable protein catalogue for future projects, as researchers can now skip directly to the targeted SRM phase to greatly reduce the cost and time associated with a discovery analysis.

Geoduck exhibited no phenotypic differences between pH conditions, counter to our predictions. We predicted that pH would be higher within eelgrass habitats, creating a refuge against the less alkaline surrounding waters and reducing oxidative stress. Concordantly, proteins involved in the oxidative stress response would be less abundant inside eelgrass habitats (such as superoxide dismutase, peroxiredoxin-1, catalase, and HSP), possibly translating to differential growth as less energy would be used to counter the pH stress. While pH in eelgrass habitats was found to be consistently higher in this study, no differences in abundance of selected peptides, growth or survival were found between habitats across all four bays. This suggests that juvenile geoduck may tolerate a wide pH range in the context of the natural environment in which they are cultured.

Earlier studies on other clam species point to some degree of pH tolerance, but also describe complex responses to low pH that vary between metrics, species, and when secondary stressors are applied (Ries et al. 2009; Ringwood and Keppler 2002; G. G. Waldbusser et al. 2010). For example, juvenile carpet shell clams (*Ruditapes decussatus*) under ambient (pH 8.2) and reduced pH (7.8, 7.5) for 75 days displayed no difference in size, weight, or calcification rate (Range et al. 2011), but other physiological parameters (clearance, ingestion, respiration, ammonia excretion) differed at day 87 (Fernández-Reiriz et al. 2011). In the hard clam *Mercenaria mercenaria*, protein oxidation, biomineralization, and standard metabolic rate (SMR, measured as resting oxygen consumption) in adults were largely unaffected by hypercapnia alone, but when combined with elevated temperature SMR increased and shell strength decreased (Ivanina et al. 2013; Matoo et al. 2013). Interestingly, the baltic clam (*Macoma balthica*) grew significantly larger in low pH (7.35 vs. 7.85 for 29 days), and were largest when combined with low dissolved oxygen (3.0 mg/L vs. 8.5 mg/L) (Jansson et al. 2015). Geoduck metrics examined in this study were not affected by varying pH, but other physiological parameters (metabolic rate, biomineralization, reproductive development, cytoskeleton), and other tissues such as mantle or hepatopancreas, may be affected and should be examined in future studies.

The complex, mixed responses exhibited in clam species may, in part, be a function of local adaptation to varying environmental drivers. Pacific geoduck are native to the Puget Sound, a region that experiences regular episodes of low pH in certain areas and has significant diel and monthly pH variability (Busch et al. 2013; Feely et al. 2008, 2010). Thus, the species may have evolved under selective pressure to withstand periods of low pH. The native Northeast Pacific Coast oyster, *Ostrea lurida*, also shows signs of pH tolerance as veliger larvae compared to the non-native Pacific oyster (*Crassostrea gigas*) (Waldbusser et al. 2016), a stage primarily found to be vulnerable in other calcifying species (for reviews see Byrne and Przeslawski 2013; Kurihara 2008). Geoduck are also infaunal organisms, extending their long siphons into the water column for feeding and retreating to deep burrows during low tide or when disturbed (Goodwin and Pease 1987). Sediment and burrow chemistry, while influenced by the overlying water column, can have lower pH due to aerobic microbial activity, another potential source of selective pressure shaping this giant clam’s pH tolerance (Gattuso and Hansson 2011; Widdicombe and Spicer 2008). An important future step is to assess the relative influence of sediment pH and overlying water column pH on burrowing calcifiers. This is particularly applicable when comparing habitats that likely have varying bacterial communities and activity.

While pH was not a universal predictor of geoduck phenotype in this study, mean temperature and dissolved oxygen variability correlated significantly with biometric parameters and separated into two groups: northern bays (Fidalgo and Port Gamble Bays), and southern bays (Case Inlet, and Willapa Bay). Geoduck grew less (or not at all, in Case Inlet) and had lower levels of targeted proteins in the southern bays, which were warmer with less variable dissolved oxygen content.

Temperatures in the southern bays (16-18°C) during the deployment dates may have exceeded optimal conditions for juvenile geoduck, resulting in elevated metabolism and less energy available for growth (Newell and Branch 1980). Similar temperature-dependent growth was observed in *M. mercenaria*, where shell calcification rate was highest between 12.8-15.2°C, above which growth negatively correlated with temperature (except for a secondary peak at 23.9°C) (Storr et al. 1982). In *P. generosa*, Goodwin (1973) reported that temperature for normal larval development is between 6-16°C. In adults, the optimal hatchery temperature for reproduction is relatively low (appr. 11°C), and at the highest experimental temperature (19°C) gonad did not regenerate after an initial spawning event (Marshall et al. 2012). Arney et al. (2015) found that in early juveniles (<3.5mm), growth increased with temperature within 7-19°C when fed ad libitum. However, organic weight accumulation (total body ash-free dry weight) was highest between 11-15°C, indicating that the optimal juvenile temperature may be approximately 15°C. In the present study, geoduck grew fastest in cooler, northern bays (15°C), but stress protein abundances (e.g. HSP90) did not suggest an acute thermal stress in the warmer, southern bays (abundances were inversely related to temperature). Southern bays may have exceeded the geoduck upper pejus temperature but remained below acute-stress, which could explain the reduced growth in those locations without a proteomic signal. Conversely, as tissues were collected at day 30, a heat stress signal could have been captured with earlier or more frequent samples. A thermal performance curve for *P. generosa* under natural feeding levels would be valuable for aquaculture siting, but these data suggest that cooler summer temperatures are more suitable for culturing geoduck.

Dissolved oxygen (DO) variability may be an indirect indicator of geoduck performance as it is often correlated to phytoplankton biomass (Khangaonkar et al. 2012). Less DO fluctuation in the southern bays could be an indicator of less phytoplankton biomass, translating to lower food availability (Anderson and Taylor 2001; Bergondo et al. 2005; Winter et al. 1975). While we were unable to monitor chlorophyll during the outplant, both southern bays, Willapa Bay and Case Inlet, may have phytoplankton populations that are controlled by shellfish grazers due to long residence times and aquaculture activity (Banas et al. 2007; Washington Sea Grant 2015). It is possible that food availability was different between northern and southern locations during the outplant period (June-July), and could be the underlying cause of higher growth and abundances of selected proteins in the northern locations (Carmichael, Shriver, and Valiela 2004; Liu et al. 2016; Loosanoff and Davis 1963), although this warrants additional data collection.

## Conclusion

This is the first study to investigate geoduck performance alongside varying pH conditions, and contributes a geoduck ctenidia peptide database useful for quantifying multiple proteins simultaneously. The primary finding is that geoduck aquaculture may be less impacted by ocean acidification compared to other environmental stressors, for example ocean warming. Geoduck ocean acidification research is in its infancy, and these results are a snapshot into geoduck physiology at one developmental stage, using one tissue type (ctenidia), with individuals from one genetic pool, and with present-day pH levels in Washington State. To best inform current and future geoduck aquaculture, further foundational studies are needed to elucidate the variability in the species’ pH limits in conjunction with more acute environmental stressors, and expanded to include other key tissues and functions (e.g. mantle for shell secretion), and whole-animal physiological studies (e.g. metabolic rate, reproductive development).

This study also demonstrates applied use of systems such as eelgrass beds in estuaries to test pH effects in a natural system. There is growing interest in using macroalgae as an ocean acidification bioremediation tool, also known as phytoremediation (Greiner et al. 2013; Hendriks et al. 2014; Washington State Blue Ribbon Panel on Ocean Acidification 2012; Groner et al. 2018). Incorporating seagrass into shellfish-pH interaction studies can help evaluate the potential for merging mariculture with shellfish aquaculture to improve growing conditions for vulnerable cultured species.

## Acknowledgements

Our gratitude to the following people who assisted with this project: Grace Crandall, Kaitlyn Mitchell and Jose Guzman assisted with protein extraction. Jarrett Egertson contributed to DIA design. Austin Keller adapted MSConvert to correctly demultiplex and convert DIA files. Han-Yin Yang, Brian Searle and Sean Bennett assisted with running PECAN. Brittany Taylor and Taylor Shellfish Farms provided geoducks. Thank you to anonymous reviewers for the helpful feedback.

This work was supported in part by the National Science Foundation Graduate Research Fellowship Program, the NOAA Ocean Acidification Program, the University of Washington's Proteomics Resource (UWPR95794), and the Washington Department of Natural Resources.

## Supplementary Material

**Supplemental Table 1.** Outplant location metrics; FB=Fidalgo Bay, PG=Port Gamble, SR=Skokomish River Delta, CI=Case Inlet, WB=Willapa Bay; -E=Eelgrass, -U=Unvegetated. ∆ Habitats, ∆ Bays, ∆ Regions: p-adjusted from 2-way ANOVA testing differences between habitats, bays, and ad-hoc regions (south=CI & WB, north=FB & PG). **Growth did not differ within ad-hoc regions. *Salinity probes failed fully or intermittently at 4 of the 8 locations; habitat salinities were not compared.

| Deployment location | FB-E | FB-U | PG-E | PG-U | CI-E | CI-U | WB-E | WB-U | Δ Habitat | Δ Bays | Δ Regions |
| --- | --- | --- | --- | --- | --- | --- | --- | --- | --- | --- | --- |
| % Survival | 87 | 80 | 80 | 93 | 67 | 60 | 93 | 100 | p= 0.88 | p=0.25 | p=0.66 |
| Mean Relative % Growth (± SD) | 16.2 (±7.3) | 23.3 (±12.5) | 22.2 (±12.5) | 21.1 (±13.3) | 1.2 (±8.3) | -2.8 (±5.3) | 10 (±16.2) | 4.5 (±8.3) | p=1 | **p= 2.4e-10 | p= 4.9e-11 |
| Mean pH (TS) | 7.90 | 7.54 | failed | 7.32 | 7.87 | 7.63 | 7.81 | 7.55 | p= 1.1e-28 | p= 1.1e-27 | p= 3.9e-3 |
| pH Standard Deviation (TS) | 0.19 | 0.23 | failed | 0.25 | 0.16 | 0.20 | 0.06 | 0.18 | p=1 | p= 1.0e-20 | p= 1.6e-15 |
| Mean DO (mg/L) | failed | 10.9 | 10.8 | 11.9 | 10.4 | 8.3 | 8.4 | 8.8 | p=1 | p= 1.1e-25 | p= 3.2e-19 |
| DO Standard Deviation (mg/L) | failed | 5.7 | 3.9 | 3.9 | 2.6 | 1.9 | 1.5 | 1.3 | p=0.31 | p= 4.6e-47 | p= 2.8e-39 |
| Mean Temp (°C) | 15.1 | 15.1 | 15.3 | 15.2 | 16.6 | 16.4 | 18.2 | 18.2 | p=1 | p= 2.2e-48 | p= 1.2e-36 |
| Temp Standard Deviation (°C) | 1.6 | 1.6 | 2.1 | 2.4 | 1.7 | 1.6 | 1.2 | 1.2 | p=0.90 | p=0.05 | p=1 |
| Mean Salinity (ppt) | 30.2 | *29.1 | 23.4 | failed | failed | 24.7 | 27.3 | *24.4 | *Not tested | *p= 2.3e-54 | p= 0.30 |
| Salinity Standard Deviation (ppt) | 0.4 | *0.34 | 1.3 | failed | failed | 1.2 | 0.70 | *1.1 | *Not tested | *p= 2.6e-3 | p=0.12 |

**Supplemental Figure 1:**
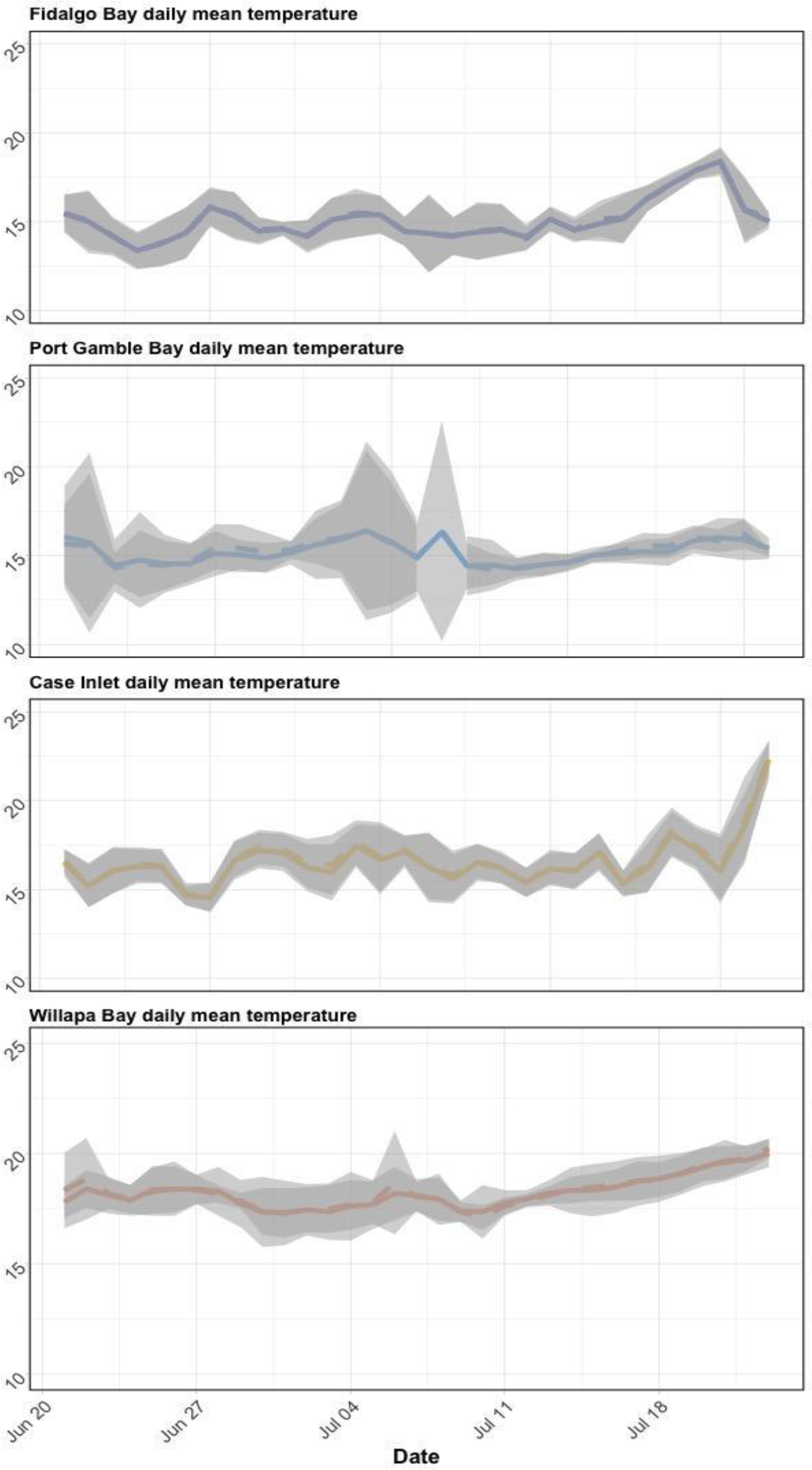
Daily mean temperature in eelgrass (dashed lines) and unvegetated (solid lines) across bays during geoduck deployment. Gray ribbons denote standard deviations per day.

**Supplemental Figure 2:**
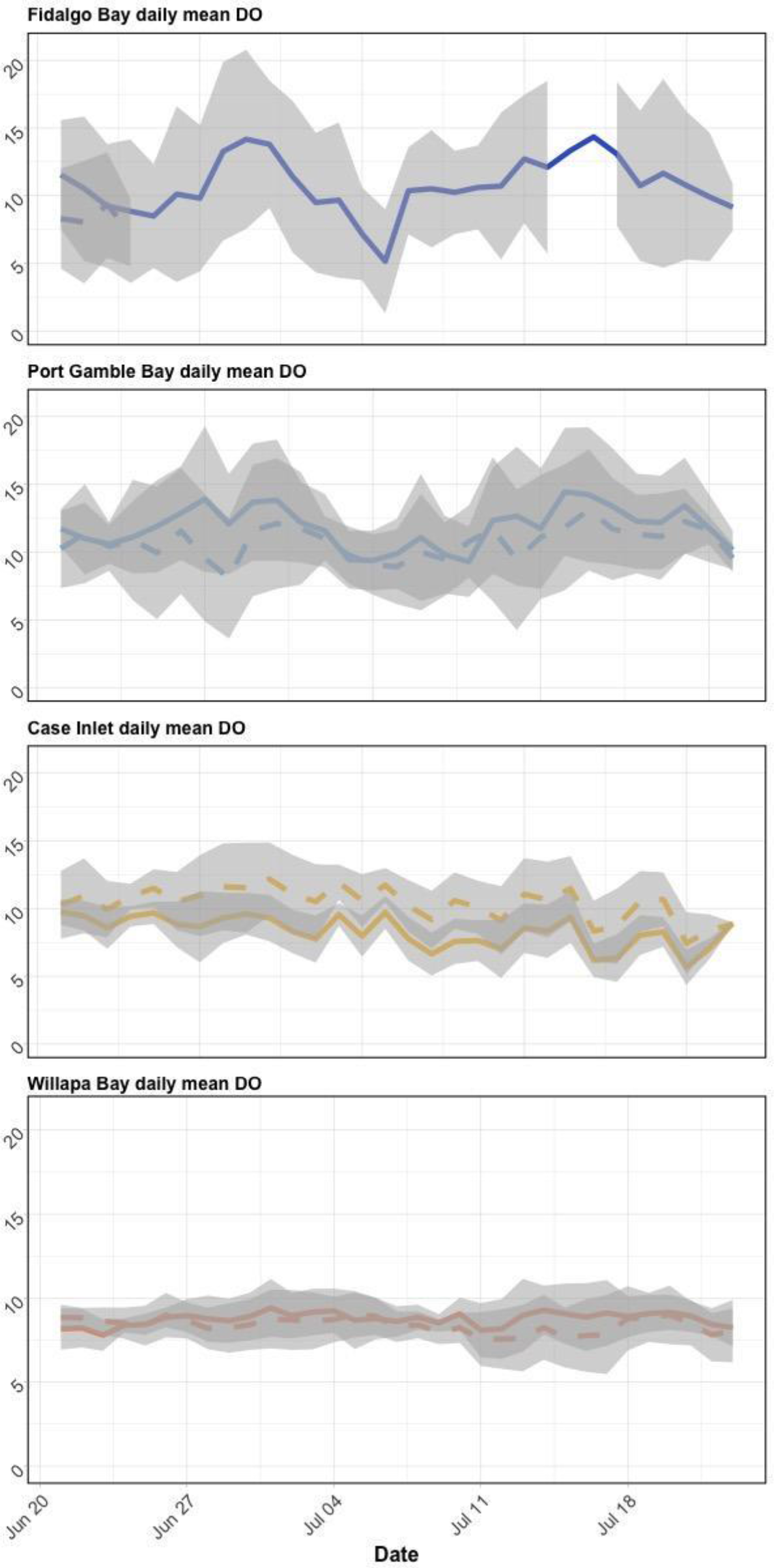
Daily mean DO in eelgrass (dashed lines) and unvegetated (solid lines) across bays during geoduck deployment. Gray ribbons denote standard deviations per day. Fidalgo Bay eelgrass probe failed towards the beginning of the outplant period.

**Supplemental Figure 3:**
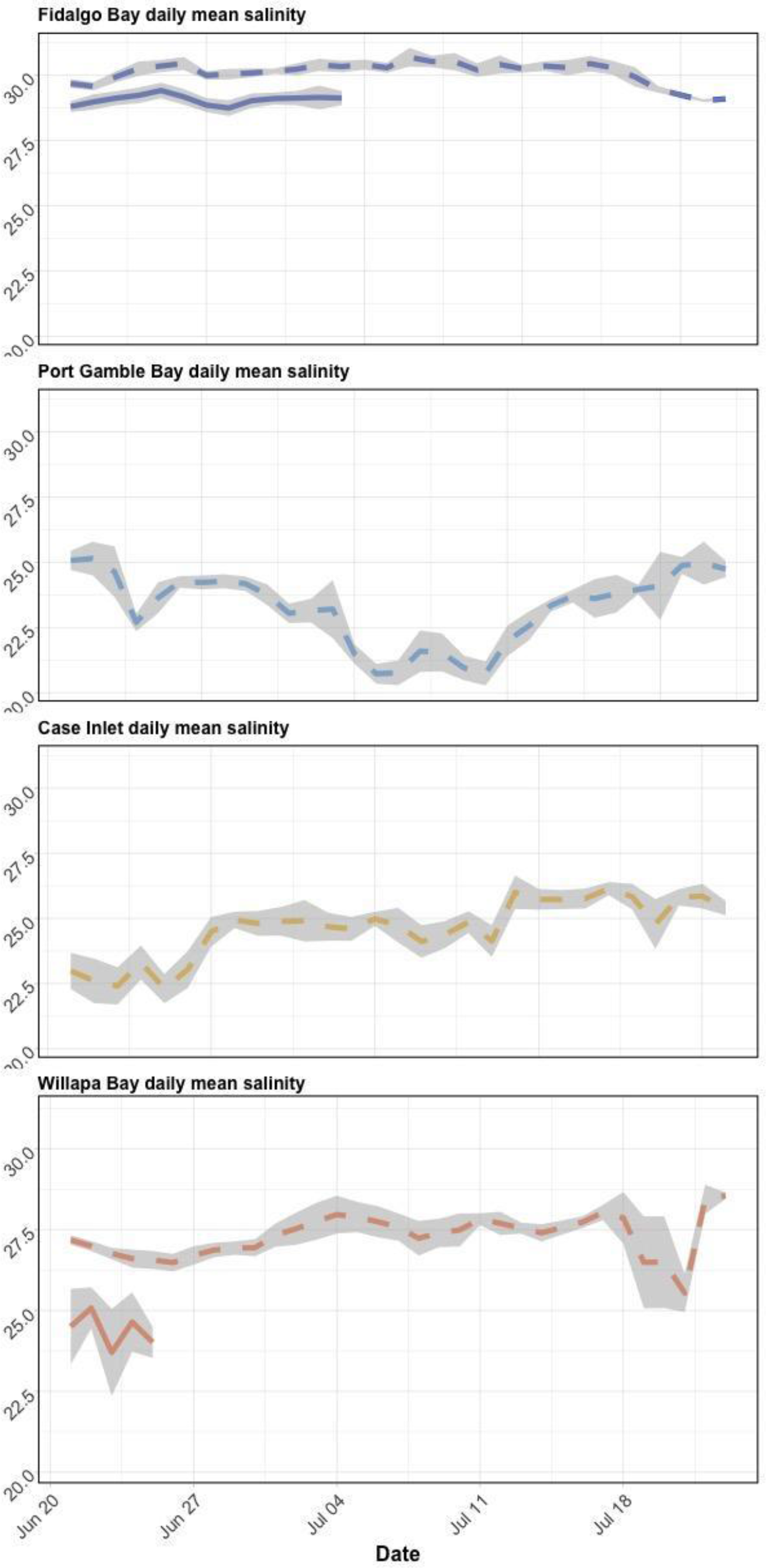
Daily mean salinity in eelgrass (dashed lines) and unvegetated (solid lines) across bays during geoduck deployment. Gray ribbons denote standard deviations per day. Salinity probes failed at several locations; no habitat comparisons were made.

**Supplemental Table 2:**
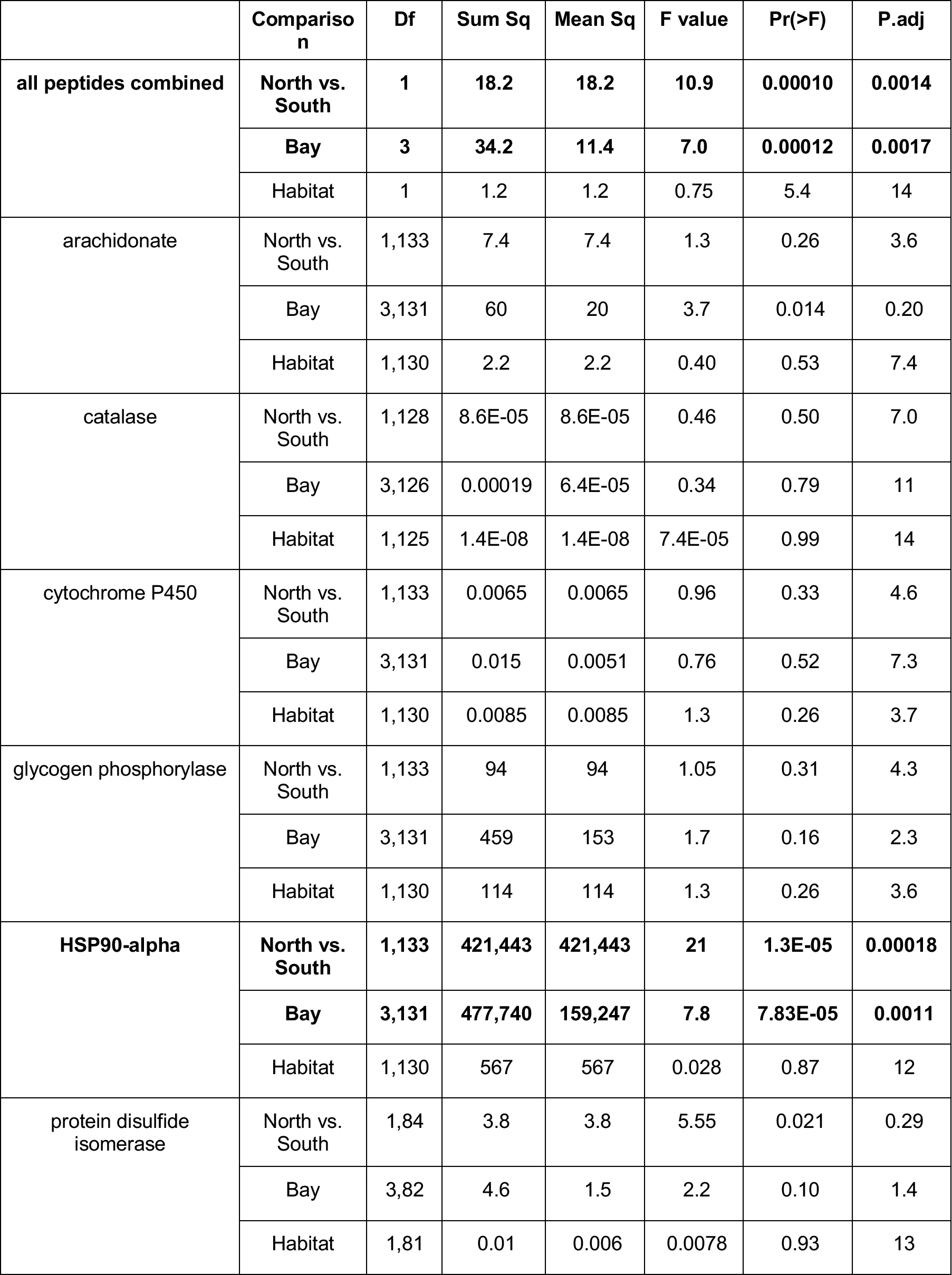
SRM protein ANOVA results by Region (FB/PGB vs. CI/WB), Bay, and Habitat for all proteins combined, then each protein individually, with Pr(>F)-adjusted calculated via the Bonferroni Correction. Bold = significantly different abundance. Habitat was tested using 2-way ANOVA with abundance ~ Bay*Habitat.

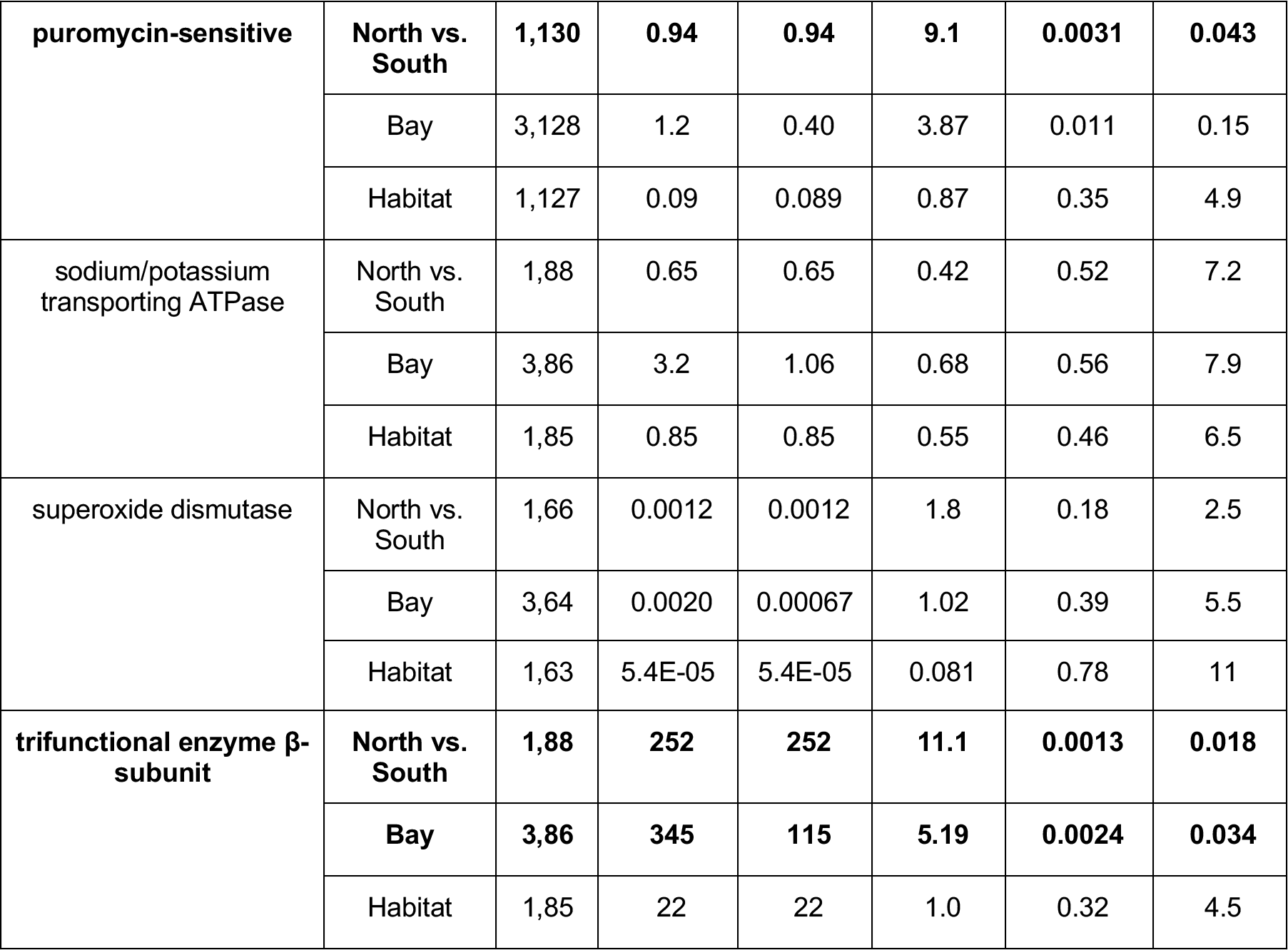

**Supplemental Table 3:**
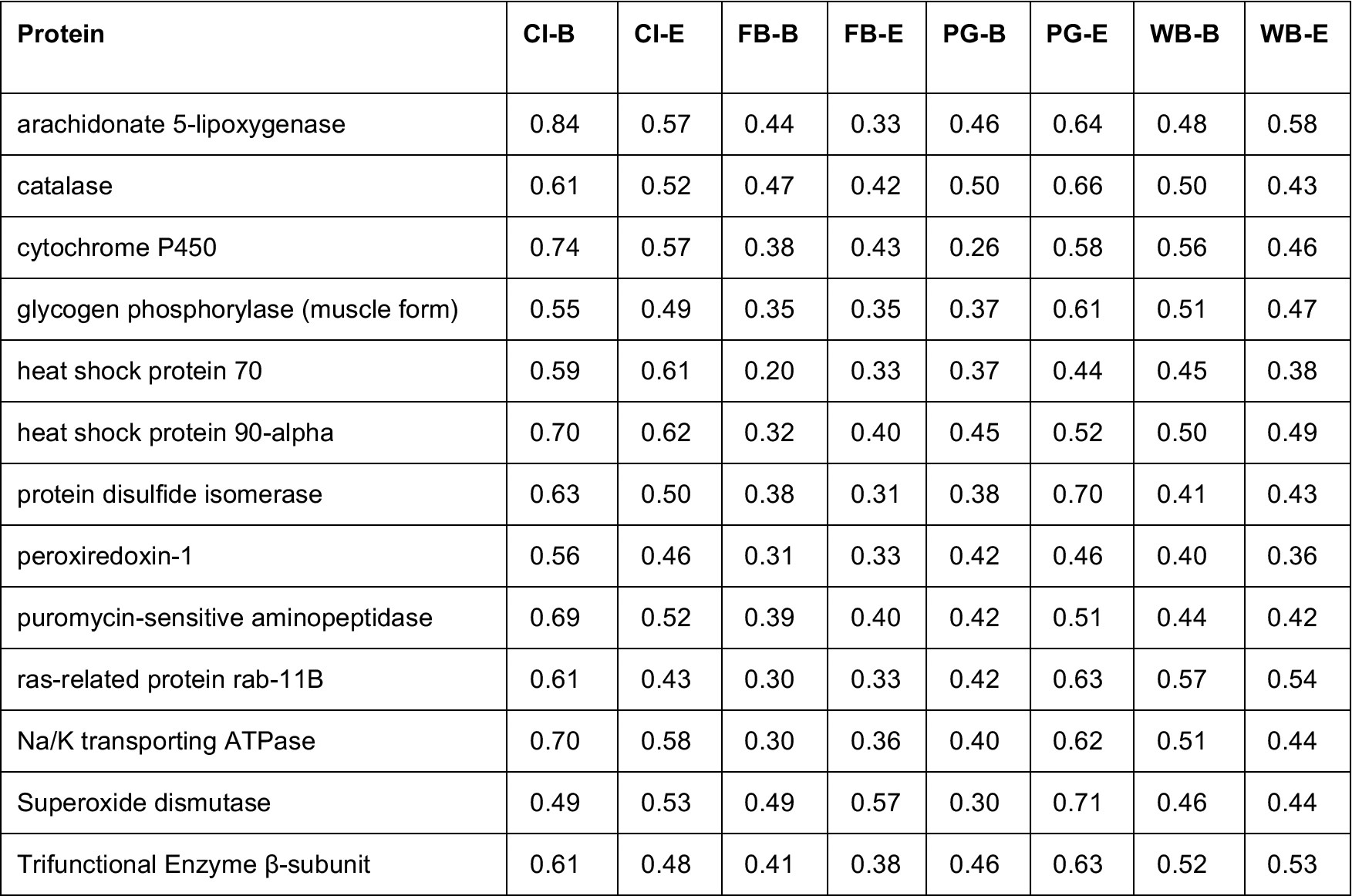
Mean coefficients of variation of SRM transition abundance for each protein, location

**Supplemental Figure 4:**
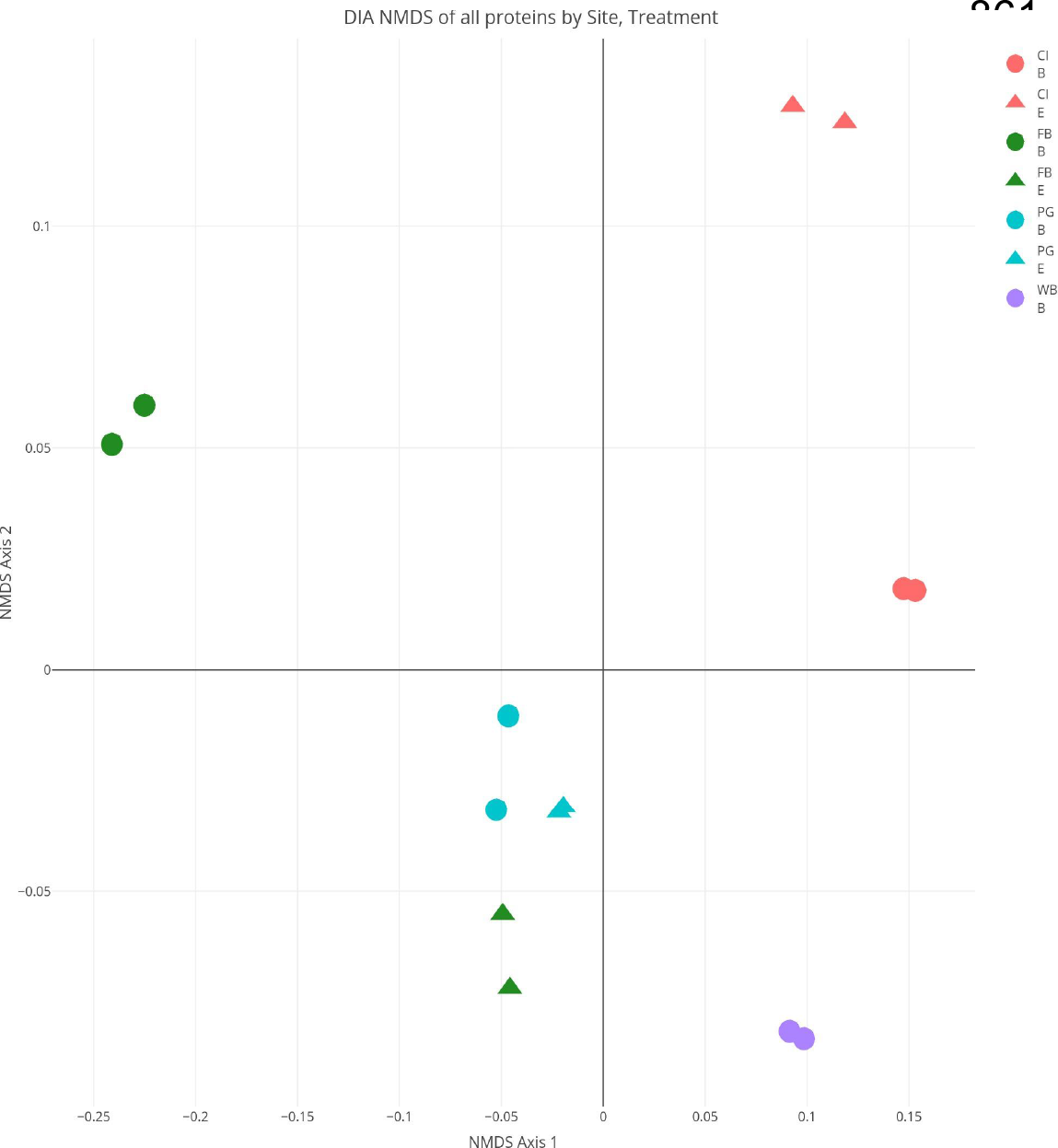
Non-metric Multi-Dimensional Scaling plot (NMDS) showing patterns of similarity among DIA peptide abundances between technical replicates (same symbol/color), deployment bay (same color), and north/south region, where each point represents one geoduck. Relative proximity of points represents overall degree of peptide abundance similarity.

**Supplemental Figure 5:**
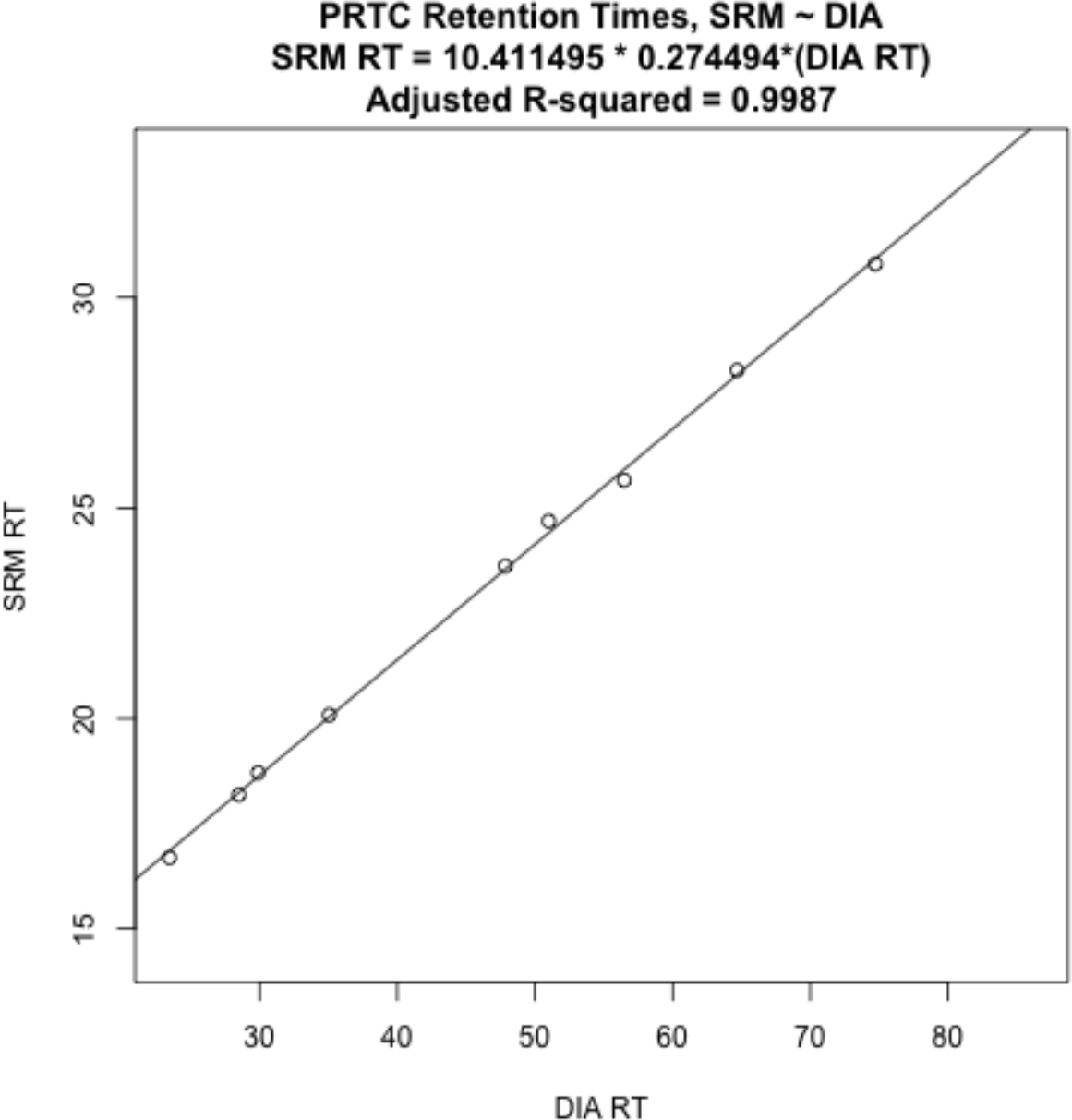
Linear regression model fit of PRTC internal standard peptides in SRM against DIA, used to confirm identity of targeted peptides in SRm by calculating adjustment in retention time between SRM and DIA.

Supplemental Figure 6 (Interactive, online):

Dilution curve peptide abundance ratios regressed against predicted ratios from serial sample dilutions. Peptides with slope coefficient 0.2<x<1.5 and adjusted R2 >0.7 were included in analysis. Link to interactive figure: http://owl.fish.washington.edu/generosa/Generosa_DNR/Dilution-Curve-Transitions.html

Supplementary Figure 7 (Interactive, online):

Non-metric Multi-Dimensional Scaling plot (NMDS) of SRM technical replicates. Link to interactive figure: http://owl.fish.washington.edu/generosa/Generosa_DNR/NMDS-technical-replicate.html

**Supplemental Figure 8:**
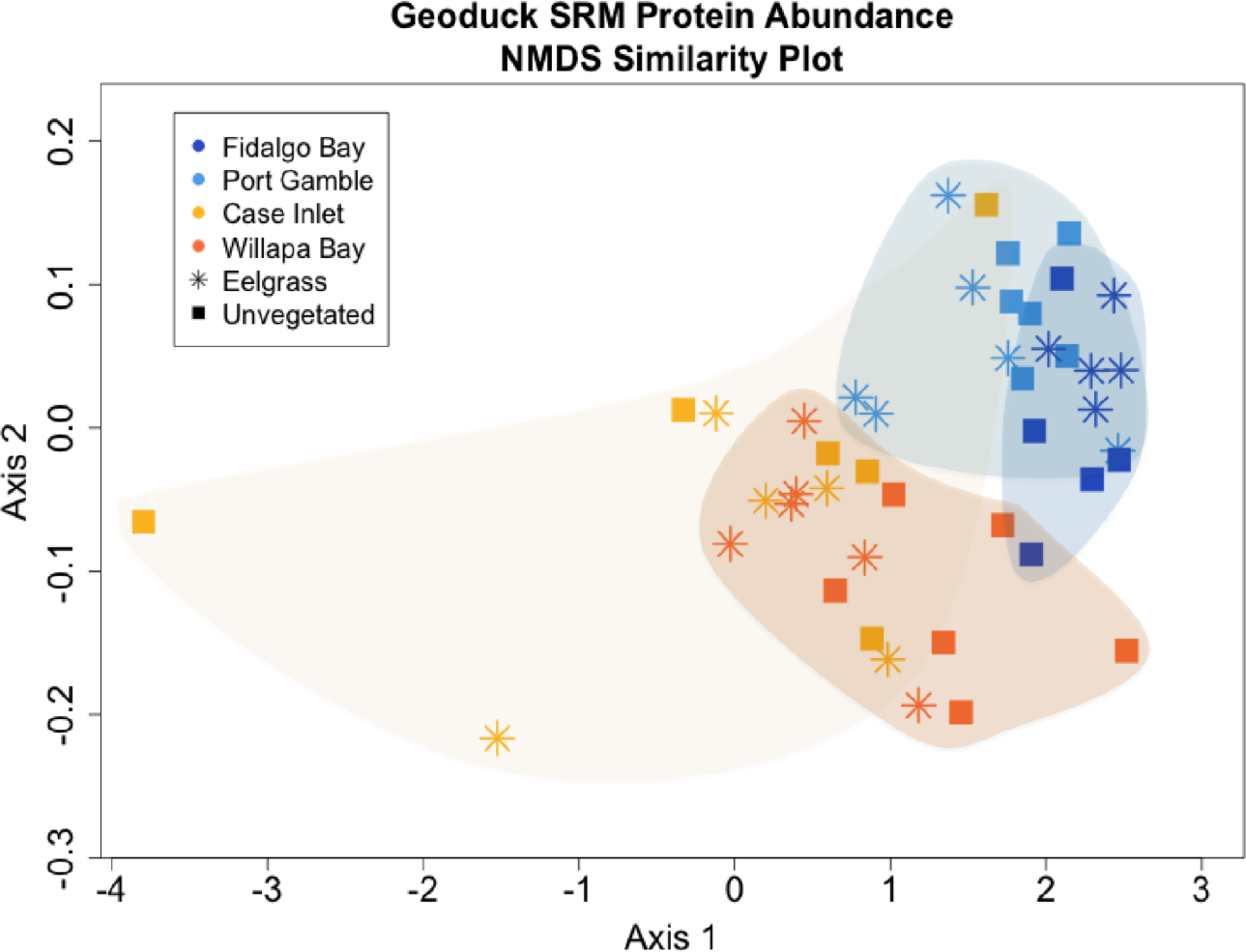
Non-metric Multi-Dimensional Scaling plot (NMDS) showing patterns of similarity among targeted SRM peptide abundances between deployment locations, where each point represents one geoduck. Relative proximity of points represents overall degree of peptide abundance similarity.

